# Establishment of menthol bleaching protocols for six stony coral species

**DOI:** 10.1101/2025.07.01.662491

**Authors:** Lavinia Bauer, Erik Ferrara, Giulia Puntin, Anna-Lena Paulus, Mia Reiser, Florian Schmidt, Marie Zeh, Maren Ziegler

## Abstract

The mutualistic symbiosis between stony corals and unicellular algae of the family Symbiodiniaceae forms the base of coral reef ecosystems. However, anthropogenic stressors, such as rising seawater temperatures, cause a breakdown of the coral-algal symbiosis, so-called coral bleaching, which leads to mass mortalities and a rapid loss of coral reefs. To functionally disassemble the coral-algal symbiosis, corals have been artificially rendered aposymbiotic using temperature stress, DCMU, or menthol. As menthol has proven to be an efficient and gentle bleaching agent, we tested four menthol treatments with six commonly investigated stony coral species. The overarching aim was to establish a broadly efficient bleaching protocol as a guide for future investigations. Menthol-induced bleaching was traced with chlorophyll fluorescence and tissue color analyses over time and confirmed as final symbiont cell density two weeks after the last day of menthol treatment. Here we found that the coral species varied greatly in their menthol bleaching tolerance, underlining the importance of establishing bespoke bleaching approaches. *Acropora muricata* and *Stylophora pistillata* were efficiently bleached within two days of menthol exposure with symbiont cell reductions of 94 % to 98 %. For *Galaxea fascicularis, Montipora digitata* and *Porites cylindrica*, six days of menthol exposure proved most successful in reducing symbiont density by 92 to 97 %. While these coral species suffered no mortality, fragments of *Pocillopora verrucosa* died or suffered severe necrosis in most of the protocols, making the tested menthol treatments unsuitable for this species. We demonstrate that repeated menthol treatment at low concentrations renders most coral species aposymbiotic within few days without visual or physiological damage. Our study, therefore, provides a guideline for efficient and customized application of menthol bleaching treatments for future coral symbiosis research.

## Introduction

The symbiosis between stony corals and the unicellular algae of the family Symbiodiniaceae forms the base for the success of highly productive coral reef ecosystems in oligotrophic waters (Frankowiak et al., 2016; Muscatine et al., 1984). However, reef-building corals and the ecosystems they build are in sharp decline all over the world. Global coral cover has declined by approximately 50 % since the 1950s, affecting not only the coral community itself but the also the biodiversity of associated organisms (Eddy et al. 2021; IPCC 2023). The increasing duration and severity of marine heat waves are the main drivers of this decline and pose an existential threat to corals worldwide (Frölicher et al., 2018). Rising sea surface temperatures cause a breakdown of the coral-algal symbiosis, so-called coral bleaching, which leads to mass mortalities that are predicted to worsen in this century (Hoegh-Guldberg et al., 2007; Hughes et al., 2018; Van Hooidonk et al., 2016).

The processes underlying coral bleaching, as well as which characteristics constitute a stress-tolerant symbiosis, remain to be fully elucidated (Berkelmans & Van Oppen, 2006; Strader & Quigley, 2022). While a benchmark study could show that heat-evolved algal symbionts may increase the heat tolerance of the coral host (Buerger et al., 2020), further mechanistic studies to interrogate the symbiotic relationship hinge on a systematic and standardized toolkit (Puntin et al., 2023). The controlled and reliable production of bleached, i.e., aposymbiotic, corals in the laboratory has therefore been a recent approach aimed at understanding the underlying processes of coral bleaching.

Methods used to date to create aposymbiotic corals include cold shock (Steen & Muscatine, 1987) and heat shock treatment (Meron et al., 2020), 3-(3,4-dichlorophenyl)-1,1-dimethylurea (DCMU) exposure (Silverstein et al., 2015), and menthol incubation ( Wang et al., 2012a). These methods pave the way to investigate the process of coral bleaching and lay the foundation to shed light on the symbiosis re-establishment process under controlled conditions. However, there is a braod range of undesirable side-effects that may hamper the downstream investigation of traits underlying heat stress tolerance or symbiosis establishment and maintenance. For instance, heat or cold treatments are time-intensive, often resulting in high mortality (Berkelmans & Van Oppen, 2006), and have strong downstream effects on the phenotype of the coral host due to environmental priming (Coffroth et al., 2010; Voolstra et al., 2020). In contrast, menthol treatment, sometimes in combination with DCMU, has the potential to efficiently produce aposymbiotic coral in small water volumes and within a few days (Scharfenstein et al., 2022).

Menthol is a cyclic terpene alcohol that acts as anesthetic on cnidarians and has first been successfully used to create aposymbiotic fragments of the coral species *Isopora palifera* and *Stylophora pistillata* (Wang et al., 2012a). Menthol bleaching did not impair host physiology and metabolism and achieved high survival rates when applied at an adequate concentration and exposure time (Wang et al., 2012a). Subsequently, menthol was also successfully used to produce aposymbiotic individuals of the model anemone *Exaiptasia diaphana* (Matthews, 2016) and the model coral *Galaxea fascicularis* (Puntin et al., 2023), and in combination with DCMU of the coral species *Diploastrea heliopora*, *Dipsastraea pallida*, *Echinopora lamellosa*, *Platygyra daedalea*, and *Porites lobata* (Scharfenstein et al., 2022). Moreover, successful re-establishment of the cnidarian-algal symbiosis after menthol treatment highlights its potential to advance coral symbiosis research (Puntin et al., 2023; Scharfenstein et al., 2022).

The efficiency of menthol to achieve complete bleaching varies between coral species and a systematic approach is needed to establish protocols for a broader range of species to enable future comparative studies. Here, we compared the effect of four menthol bleaching protocols on six stony coral species to identify the protocol with highest survival and bleaching efficiency for each species. As these represent the most-commonly used species in bleaching studies (McLachlan et al., 2020), our work provides a tool for future coral symbiosis research.

## Material and Methods

### Experimental design

The bleaching experiment was conducted at the *Ocean2100* aquarium facility at Justus Liebig University Giessen, Germany. Long-term rearing conditions were a light:dark cycle of 12:12 h at 174-263 µmol photons m^-2^ s^-1^ supplied by white and blue fluorescent lamps, salinity of 36, and temperature of 26.5 °C. Coral colonies were fed daily with a mixture of frozen copepods, Artemia, krill, and Mysis (Schubert & Wilke, 2018).

We used six common reef-building coral species: *Acropora muricata* (Linnaeus, 1758)*, Stylophora pistillata* (Esper, 1792), *Pocillopora verrucosa* (Ellis & Solander, 1786)*, Galaxea fascicularis* (Linnaeus, 1767)*, Montipora digitata* (Dana, 1846), and *Porites cylindrica* (Dana, 1846) in two experimental runs (Tab. S1). Run 1 was carried out in 2022 and included the treatments 1m, 1m6r1m, and 3m1r1m for all six species. Early results showed that *G. fascicularis, M. digitata,* and *P. cylindrica* did not bleach sufficiently with these bleaching protocols. Consequently, run 2 was carried out in 2023, and included a fourth menthol treatment specifically for these three species with additional treatment days (3m4r3m). Three source colonies per species were selected, and from each colony, eight fragments of 3-5 cm were produced for run 1, and four fragments for run 2 because of the lower number of treatments in that run. Fragments were suspended on nylon strings and allowed to recover for 3-5 weeks before the start of the experiment. Fragments were randomly distributed among the treatments (n = 6 per treatment per species), which included a control that was not exposed to menthol and four distinct menthol treatment protocols.

### Menthol bleaching protocols

To establish the most effective bleaching protocol for each species, we exposed six reef-building corals species to four menthol treatments. Initially, three treatments were tested (run 1) while the fourth treatment was later added for *G. fascicularis*, *M. digitata*, and *P. cylindrica* due to incomplete bleaching results obtained with the first run. Coral fragments were incubated in 1-L glass containers filled with 65-µm sieved seawater from the aquarium system. Each jar contained three (menthol) or five (control) coral fragments. The seawater was spiked with menthol (stock solution: 1.28 M, 20 % m/v menthol in 99% ethanol) and lids were loosely fit to minimize evaporation. Coral fragments were incubated for 8 h per day at 26 °C, corresponding to the long-term rearing temperature, under constant illumination (200 ± 10 µmol photons m^-2^ s^-1^) and stirring (∼2.62 cm/s) (Rades et al., 2022). The menthol concentration and the number of days of incubation varied between treatment: 1) **1m**: one day of menthol incubation at a menthol concentration of 0.58 mM. Due to the strong negative response of *P. verrucosa* to menthol after the incubation, we lowered the menthol concentration to 0.19 mM in the 1m treatment and in the second day of 1m6r1m treatment for this species; 2) **1m6r1m:** one day of <u>m</u>enthol incubation, followed by 6 days of <u>r</u>est and a second day of incubation at a <u>m</u>enthol concentration of 0.58 mM.; 3) **3m1r1m**: three days of menthol incubation, followed by one day of rest, and a fourth incubation day at a menthol concentration of 0.38 mM. This treatment is an adaptation from the protocol by Wang et al. (2012) and effectively produced fully bleached single polyps of *G. fascicularis* in a previous experiment (Puntin et al., 2023); 4) **3m4r3m**: three days of menthol incubation, followed by four days of rest, and another three days of incubation at a menthol concentration of 0.38 mM. The 3m4r3m treatment was included only in run 2 exclusively for *G. fascicularis, M. digitata*, and *P. cylindrica*. In the control coral fragments were incubated following the 3m1r1m (run 1) or 3m4r3m (run 2) protocol in menthol-free seawater to account for the potential stress induced by the incubation procedure.

At the end of the 8-h incubations, each fragment was consecutively dipped into three beakers filled with menthol-free seawater to rinse off the menthol and placed back in the respective holding tank. Coral fragments were fed every other day with frozen copepods (concentration of 3-4 mg L^-1^), but not on menthol incubation days.

### Bleaching assessment

Bleaching was assessed at specific time points relative to beginning and end of the menthol treatment. The timepoints included the day before the first menthol incubation and two, seven and 15 (run 1)/ 13 (run2) days after the last menthol incubation (Fig.1, Fig. S1). These timepoints (-1, 2, 7, 13 or 15 days after menthol) are referred to as t0, t1, t2, t3, respectively.

**Fig. 1:**
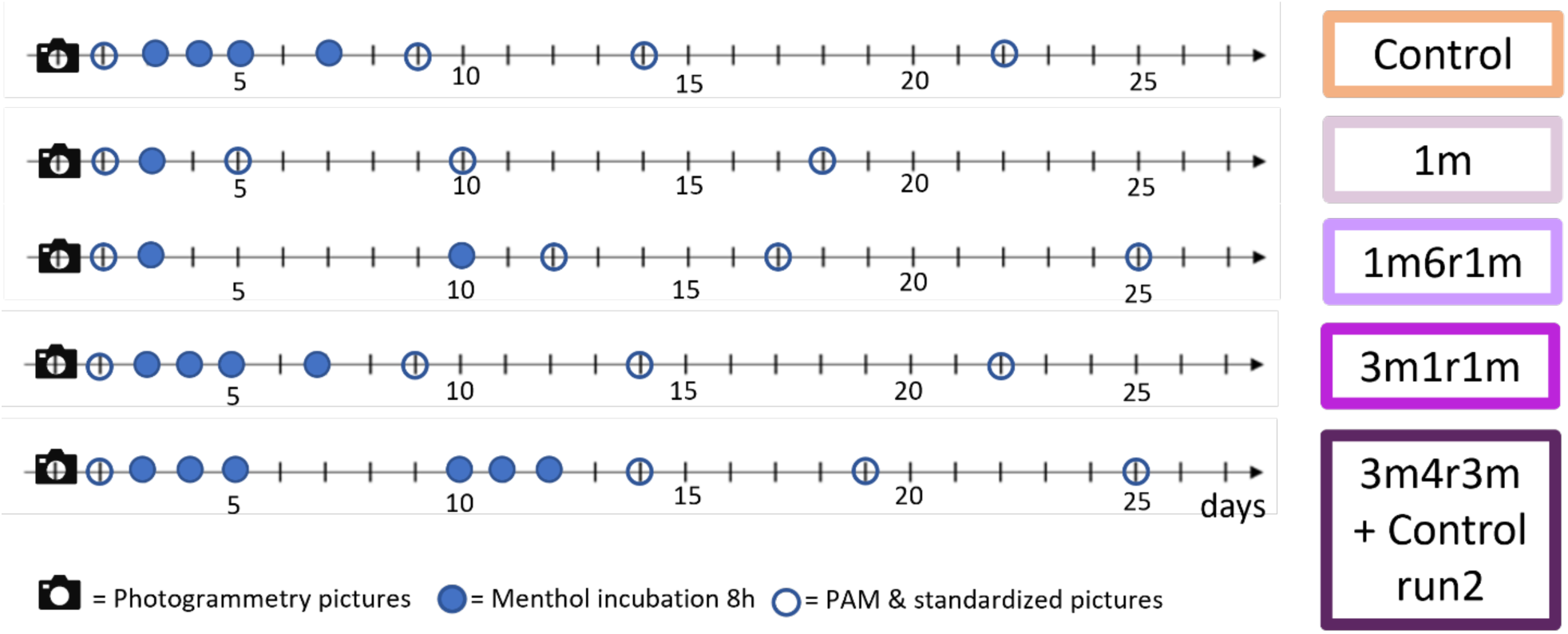
Timelines of the menthol bleaching treatments applied to six stony coral species. Each tick mark on the axis represents one day. Filled circles show days with menthol incubations (8 h per day), empty circles show days with measurements of photosynthetic efficiency and acquisition of standardized pictures, the camera symbol indicates the day pictures for surface area calculations were taken.

We assessed the effectiveness of the four bleaching treatments by repeated measurements of light-adapted minimal fluorescence (F_0_) and tissue color intensity at t0-t3, and determination of symbiont cell density at t3.

### Light-adapted minimal chlorophyll fluorescence

Minimal fluorescence of chlorophyll (F_0_) was measured with a Pulse-Amplitude-Modulated fluorometer (PAM-2500, Walz, Germany) fitted with the manufacturer’s distance clip to keep a 45° angle between the sample plane and the fiber optic. At each timepoint, each fragment was measured at three different positions around noon using the default settings (Measuring Light int.=4, MF.L=200, MF.H=20000;, Red Actinic Light int.=1, Gain=1, Sat.Pulse int.=8, Sp-width=8). The mean of the three values was computed and used for further analysis. Because of the severe bleaching, photosynthetic efficiency could not be reliably quantified and we used F_0_ as measure of chlorophyll content.

### Tissue color intensity

To assess the degree of menthol-induced bleaching, we extracted color values from standardized pictures of the coral fragments. They were documented with a DSLR camera (Nikon D7000 with a Tamron 90mm macro lens) in an evenly illuminated soft box over a black background. Orientation of fragments was maintained consistent between time points, and the same settings were used for all pictures (aperture: F10, shutter speed: 1/20, ISO: 400, and white balance: 5560K). The pictures were then processed with Adobe Photoshop CS6 and/or Meta AI Segment Anything (Kirillov et al., 2023) for white balance and to remove the background. The pictures were then saved with a transparent background in PNG24-format. Values for each color channel (red, green, and blue) and grayscale conversion were extracted and calculated with a custom Python script (Reichert et al., 2021). We have previously shown that relative changes in coral color scale linearly with changes in symbiont densities, with the red channel as best proxy for most species (Ferrara et al., 2024) and we therefore focus on the values measured from red channel herein.

### Photogrammetry for surface area calculations

Coral fragment surface area was calculated from photogrammetry. Approximately 50 pictures of each fragment were taken 2 days before the respective first menthol incubation. The pictures were processed to create 3D models in 3DF Zephyr Free (v.6.507 / 6.513/ 7.003). Model cleaning was done with Artec3D (Studio 16 Professional v.16.0.8.2). Size scaling and calculation of live coral surface area of cleaned models was done in MeshLab (v.2016.12, (Cignoni et al., 2008)).

### Symbiont count

The coral fragments were frozen at -20 °C after the last measurements at t3. Symbiont cells were extracted in 50-mL Falcon tubes using 1M NaOH (Zamoum & Furla, 2012). Tubes were placed in a 36 °C water bath for 60 minutes and shaken every 15 minutes. The clean coral skeleton was removed from the tube, and the suspension was centrifuged at 3,000 g for 5 min (Heraeus Instruments, Laborfuge 400R). The supernatant was removed, and centrifugation was repeated for 1-2 wash steps in 10 mL 0.22-µm filtered seawater. Symbiont cells were counted at 400× magnification using a Thoma cell counting chamber (Marienfeld, DIN 12847). Six chambers were counted per sample and symbiont cell density was normalized to the surface area of the coral fragments.

### Statistical analyses

Data analyses were conducted in R (4.1.3) and RStudio (2023.06.0) (R Core Team 2021). For minimum fluorescence and tissue brightness, the difference between menthol treatments (1m, 1m6r1m, 3m1r1m, 3m4r3m were analyzed separately per species using one-factorial-repeated-measures ANOVA followed by post-hoc paired t-tests per sampling time point (t0-t3) (Kassambara 2023). Bonferroni-correction was applied for the post-hoc comparisons to correct for multiple testing. Symbiont density at the end of the menthol treatments (t3) was analyzed with a one-factorial-ANOVA followed by a post-hoc Tukey-HSD-test for pairwise comparison. All plots were generated with ggplot2 (Wickham 2016).

## Results

Testing four different menthol treatments on six stony coral species allowed us to identify the most efficient bleaching treatment per species, as coral species showed high variability in menthol bleaching susceptibility. In all species but *P. verrucosa,* all coral fragments survived the menthol bleaching treatments. *P. verrucosa* was highly sensitive to menthol, with a 100 % mortality rate of fragments exposed to the 3m1r1m treatment and to initially high menthol concentrations. Additionally, *P. verrucosa* fragments exhibited necrotic tissue after menthol exposure in 1m6r1m (100 %) and 1m (67 %) despite the lower concentrations used in this treatment (Tab. S2). Therefore, due to the lack of data and statistical power, *P. verrucosa* was excluded from statistical analyses, and it was therefore not possible to identify a suitable bleaching protocol.

### Minimal chlorophyll fluorescence

Minimal fluorescence (F_0_) of the chlorophyll present in *Symbiodiniaceae* was measured as proxy of algal symbiont presence. Minimal fluorescence was significantly affected by the menthol treatments, and the interaction of treatment and time for each species (Two-Way ANOVA, p ≤ 0.05) (Tab. S3). For all menthol treatments, F_0_ was lowest at day 2 after menthol incubation (t1) and subsequently increased by 12-97 % by the end of the measurement period (Fig. 2a).

**Fig. 2:**
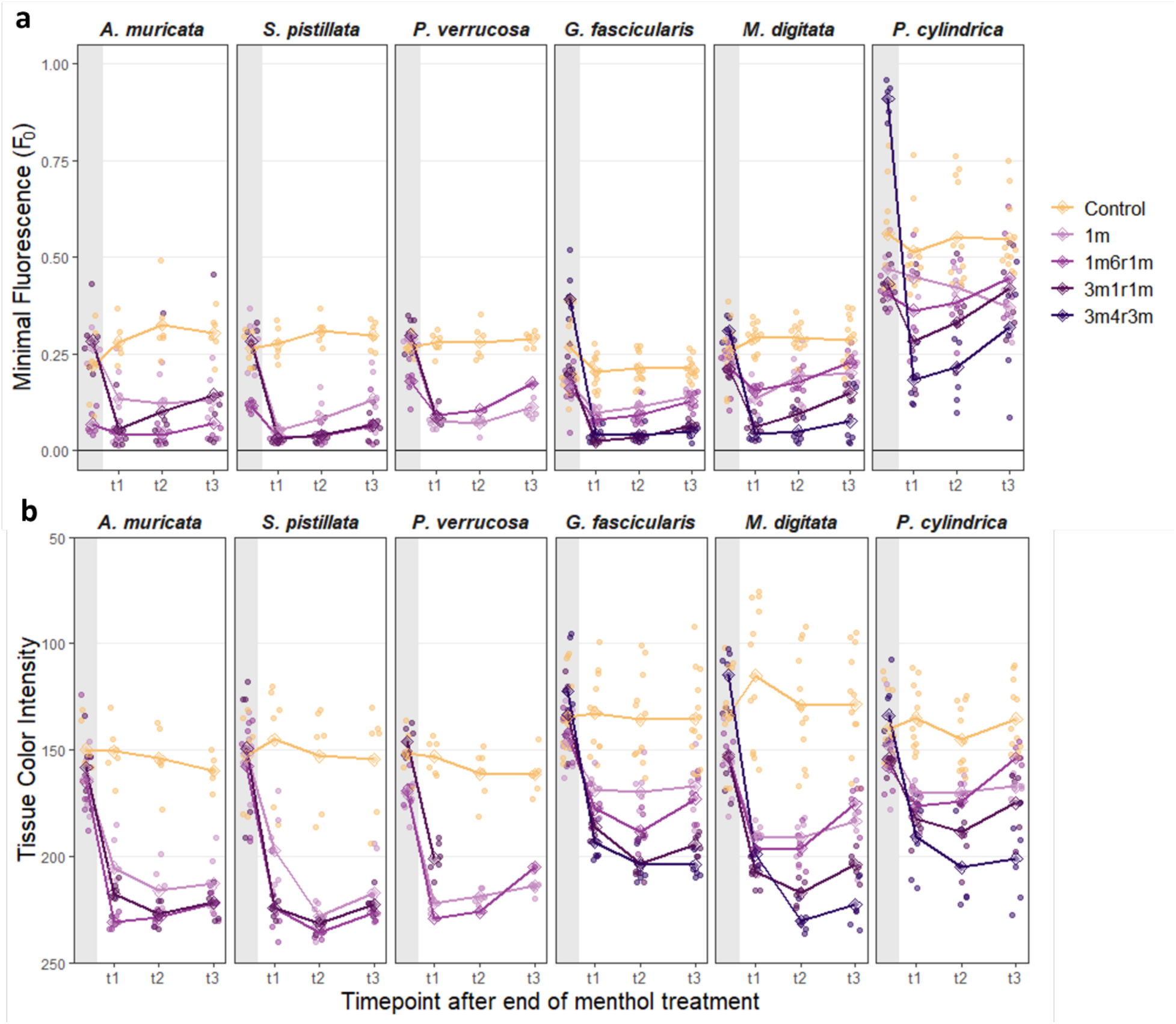
Effect of four menthol bleaching treatments on symbionts of six stony coral species. a) Light-adapted minimal chlorophyll fluorescence (F_0_) and b) tissue color intensity at the time points t0 before (gray shading) and t1, t2, and t3 after menthol treatments. For tissue color intensity, 0 corresponds to black and 255 to white (note reversed y-axis). Line colors represent different treatments with darker purple corresponding to longer menthol exposure. Diamonds represent mean values of each treatment per timepoint.

Minimum fluorescence was significantly lower in all menthol treatments than in the control at time points t1 and t2 for all species (Tab. S4) except for *P. cylindrica* which did not differ from the control in the 1m treatment (p > 0.05). At time point t3, F_0_ in *M. digitata* and *A. muricata* had increased again, reaching levels that were not statistically different from the control in the 1m6r1m (p > 0.05) and the 3m1r1m treatment (p > 0.05). F_0_ was similar among all menthol treatments in *A. muricata* and *S. pistillata* (p > 0.05). Additionally, no significant difference in F_0_ was observed between the most intense treatment, 3m4r3m, and the 3m1r1m treatment in *G. fascicularis*, *M. digitata*, and *P. cylindrica* (p > 0.05; Tab. S4).

### Tissue color intensity

To trace bleaching patterns across treatments over time, we measured tissue color intensity by extracting the red, blue, green, and gray channel values from standardized, white-balanced pictures (Fig. 3). Here, we present data from the red channel only, as it resolved the fine-scale color intensity differences better than the other channels (Ferrara et al. 2024; Fig. S4).

**Fig. 3:**
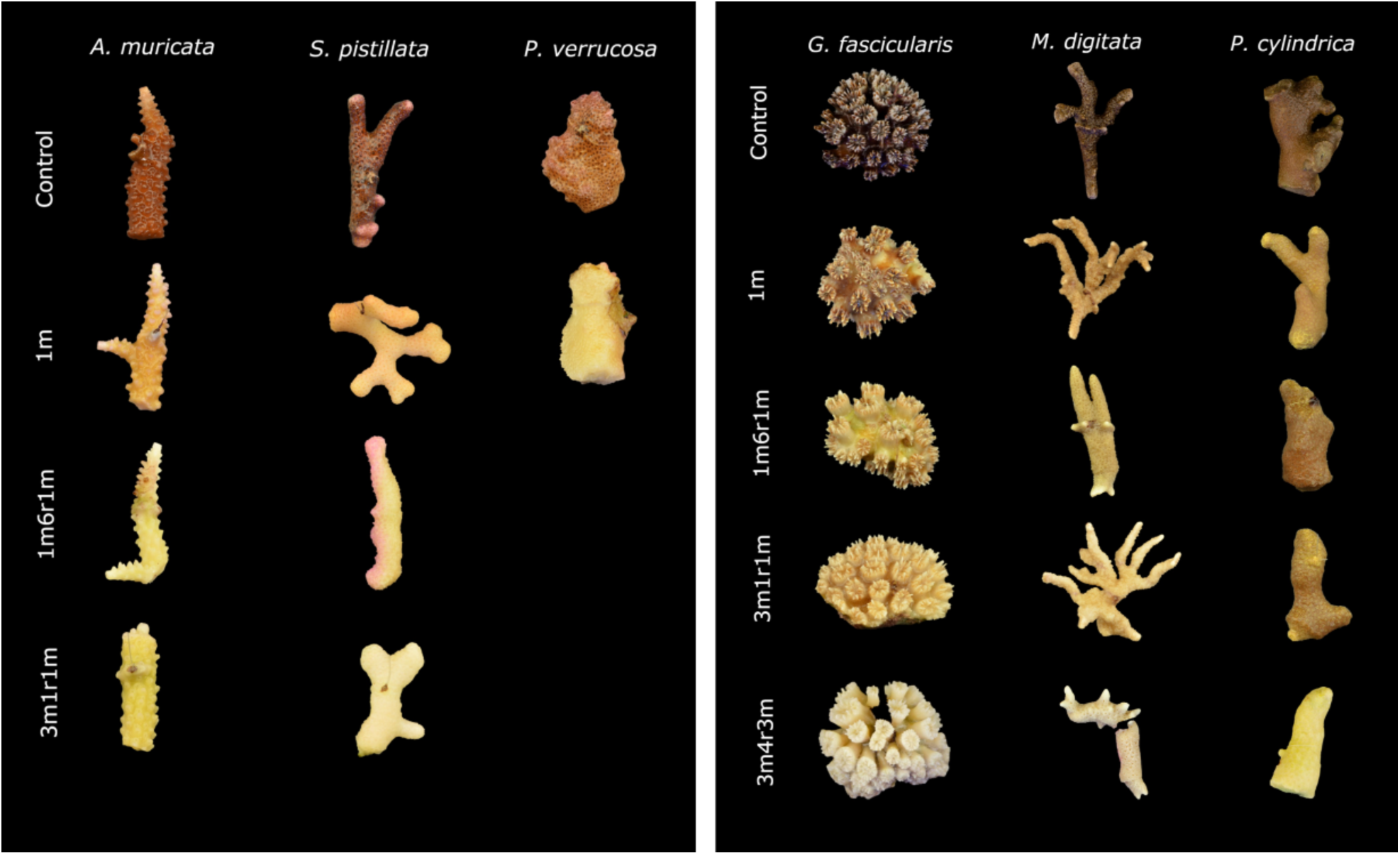
Coral fragments at sampling time point t3 after different menthol bleaching treatments. Representative fragments of different genotypes with standardized pictures of *A. muricata, S. pistillata, P. verrucosa, G. fascicularis*, *M. digitata*, and *P. cylindrica*. *P. verrucosa* died in the treatments 1m6r1m and 3m1r1m (detailed bleaching series Fig. S3).

The effect of treatment, time, and the interaction of treatment and time on bleaching severity was significant across all species without exception (p ≤ 0.001, two-way repeated measures ANOVA, Fig. 3, Tab. S5). At t1-t3 after the menthol treatments, coral fragments had bleached, represented by a significant decrease in tissue color intensity across all treatments and species compared to the control (Fig. 2b, Tab. S5, S6). Overall, treatments 3m1r1m and 3m4r3m caused more pronounced bleaching (lighter tissue color) compared to the 1m, 1m6r1m, and control treatments (p ≤ 0.05, Tab. S6).

For *A. muricata,* bleaching severity significantly increased between the single (1m) to double menthol treatment (1m6r1m), but not when additional treatment days were added (1m6r1m to 3m1r1m) (Fig. 2b, Tab. S6). In *S. pistillata*, all coral fragments bleached similarly in all menthol treatments, which were all significantly brighter than the control fragments (Fig. 2b, Tab. S6). For *P. cylindrica*, *M. digitata*, and *G. fascicularis*, the 3m4r3m menthol treatment resulted in highest bleaching severity (Fig. 2b, Tab S6). *P. cylindrica* exhibited a significant decrease in tissue color intensity in the 3m4r3m treatment compared to the 3m1r1m treatment at t3 (p < 0.05, post-hoc paired t-test). While this trend was similar for *G. fascicularis* and *M. digitata*, no significant differences were observed between 3m4r3m and the 3m1r1m treatments at the level of tissue color intensity (p > 0.05, post-hoc paired t-test).

### Symbiont cell density

Symbiont cell density at the end of the experiment was significantly lower in all menthol treatments compared to the control for all coral species (Fig. 4, Tab. S7). In *A. muricata* and *S. pistillata* symbiont density in the 1m6r1m and 3m1r1m treatment was reduced by 93 and 98%, respectively, compared to the control (Fig. 4). Symbiont density in the 1m treatment was slightly higher (reduced by 73-80 %), but still within a range with the other treatments (1m:1m6r1m, p=0.393; 1m:3m1r1m, p=0.419; Tukey HSD for *S. pistillata*), which were all significantly lower than the Control (p ≤ 0.001, Tukey HSD) (Fig. 4, Tab. S7).

**Fig. 4:**
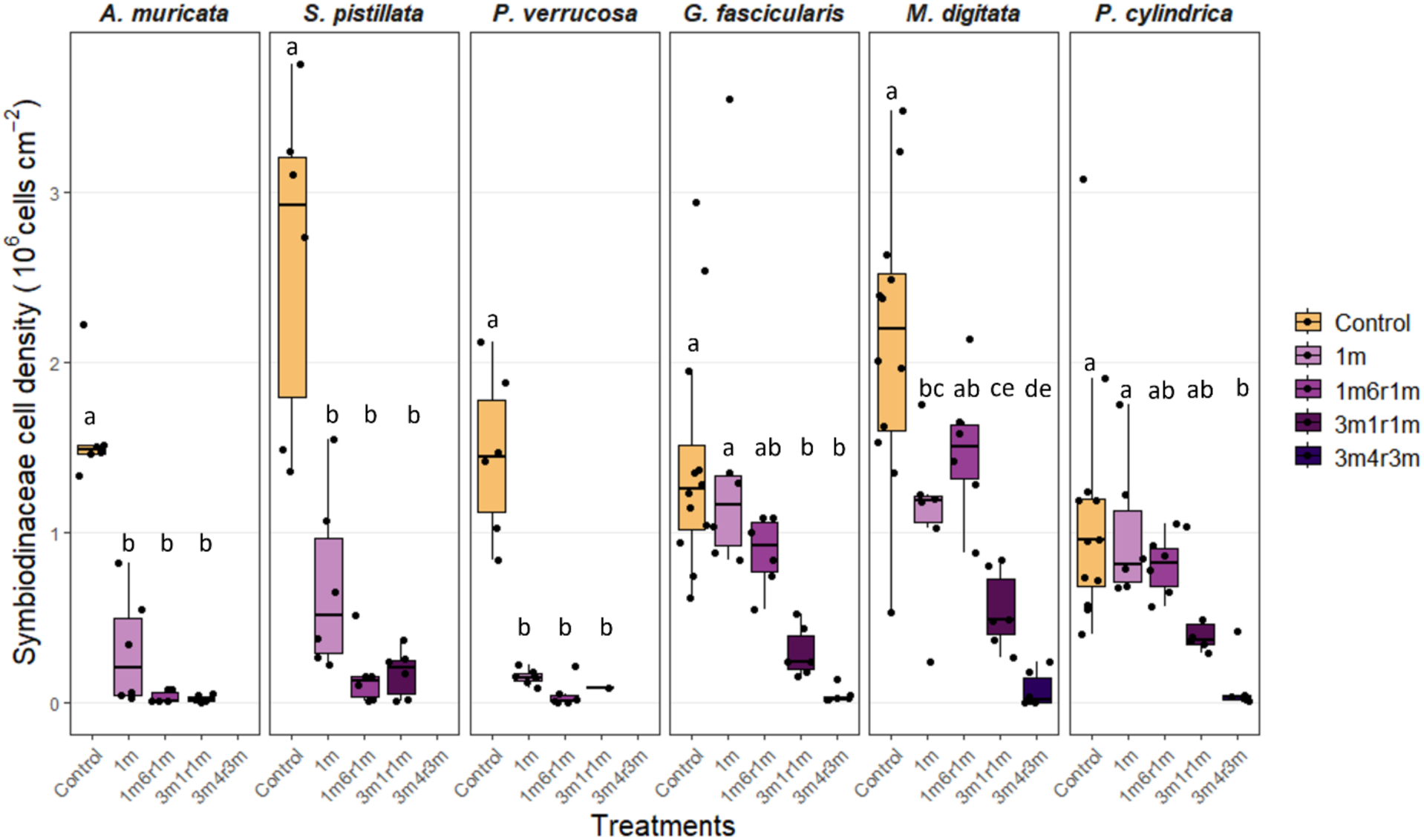
Symbiont cell density of six coral species at the end of a menthol bleaching experiment. Colors show different menthol treatments – darker purple shade indicates increased menthol exposure; letters (a, b, c) indicate significant differences (one-way ANOVA, Tukey’s-HSD post-hoc test). Data are displayed as box-and-whisker plots with raw data points; lines indicate medians, boxes indicate the first and third quartile, and whiskers indicate ± 1.5 IQR.

In *G. fascicularis*, *M. digitata,* and *P. cylindrica* symbiont density was lowest at the end of the 3m4r3m treatment, with only 45,000, 78,000, and 92,000 cells cm^-2^ remaining, respectively, corresponding to a reduction of 92-97 %. The 3m1r1m treatment had intermediate symbiont densities compared to the Control and treatments 1m and 3m4r3m (Fig. 4). Symbiont density in the 1m and 1m6r1m treatments were only slightly decreased (12-38 %) with respect to the Control in these species (p ≥ 0.1, Tukey HSD; Fig. 4, Tab. S7). The high mortality in *P. verrucosa* resulted in low sample replication, which hampered a detailed comparison between treatments, which all had significantly lower symbiont densities than the Control (p ≤ 0.002, Tukey-HSD; Fig. 4, Tab. S7).

## Discussion

The aim of this study was to establish the menthol bleaching protocol with highest survival and bleaching efficiency for each coral species. We showed that the coral species reacted differently to menthol exposure, with each species requiring a bespoke protocol to optimize bleaching. For instance, while *P. cylindrica* exhibited effective bleaching only in response to the most intense treatment, *P. verrucosa* was very sensitive and suffered high mortality after menthol treatment. Based on our data, we propose specific bleaching protocols for six coral species, that will enable future symbiosis research (Tab. 1).

**Tab. 1:** Recommendation for optimized coral species-specific menthol bleaching protocols. Menthol bleaching protocols are shown by name including respective menthol concentrations. Abbreviations consist of the number of m =menthol incubation days and r = rest days according to the protocol sequence. For *P. verrucosa* menthol concentration in 1m and 1m6r1m was lowered to 0.19 mM.

| *Species* | *Menthol bleaching protocol* |  |  |  |
| --- | --- | --- | --- | --- |
|  | 1m<br>(0.58 mM) | 1m6r1m<br>(0.58 mM) | 3m1r1m<br>(0.38 mM) | 3m4r3m<br>(0.38 mM) |
| *Acropora muricata* |  | x |  |  |
| *Stylophora pistillata* |  | x |  |  |
| *Pocillopora verrucosa* | x<br>(0.19 mM) |  |  |  |
| *Galaxea fascicularis* |  |  |  | x |
| *Montipora digitata* |  |  |  | x |
| *Porites cylindrica* |  |  |  | x |

### Species-specific treatment efficiency

The six investigated species could be divided in two groups based on menthol bleaching tolerance. *A. muricata, S. pistillata,* and *P. verrucosa,* were sensitive, while *G. fascicularis, M. digitata,* and *P. cylindrica* were tolerant to menthol exposure. Such differences in bleaching tolerance between species have previously been reported. Specifically, the bleaching tolerance of the genus *Porites* observed herein aligns with the work of Scharfenstein et al. (2022), who reported that *Porites lobata*, a close relative of *P. cylindrica*, retained more symbionts after menthol exposure than any other species. Accordingly, we developed a fourth, more intense treatment to achieve near-complete bleaching in this species. *M. digitata* behaved similarly to *P. cylindrica* in our study, despite its distance in phylogeny and other traits (Madin et al., 2016). *A. muricata* and *S. pistillata* reached maximum levels of bleaching in the mildest menthol treatments (1m, 1m6r1m), while further increase in menthol exposure did not induce signs of physiological damage. Prior studies have achieved successful bleaching in *S. pistillata* (Scharfenstein et al., 2022; Wang et al., 2012a), which was confirmed by our observations. *S. pistillata* bleached rapidly and efficiently within only two menthol incubations and hence is a promising species for further symbiosis manipulations. Other species, including the coral model *G. fascicularis* (Puntin et al., 2023) required higher doses or longer exposures to menthol, but this does not preclude them from subsequent, successful symbiosis manipulation (Chan et al. 2023).

Notably, the sensitivity to menthol-induced bleaching was not relatedness of the coral species at the family level. Within the family Acroporidae, *A. muricata* and *M. digitata,* and within the Pocilloporidae, *P. verrucosa* and *S. pistillata* each showed clear differences in their tolerance to menthol bleaching. These findings suggest that coral phylogeny might not be a reliable predictor for the menthol bleaching response. Specifically, the high mortality in *P. verrucosa* might have been affected by other rearing and handling factors such as the feeding regime which might be crucial in maintaining healthy bleached coral as their primary energy source is lost during bleaching (Muscatine et al., 1984). The coral fragments in this study were fed every other to every day. However, we did not supply additional nutrients that have been shown to potentially improve coral health (Wang et al., 2012a; Lopez et al., in preparation). Thus, to increase the probability of coral health and survival for e.g., further re-inoculation experiments, it might be beneficial to implement additional supplementation during menthol bleaching approaches.

Differences in the bleaching response may be explained by differences in Symbiodiniaceae associated with the coral. Symbiodiniaceae communities vary across species, including differences in relative abundance of dominant and potentially several other background symbionts (Berkelmans & Van Oppen, 2006; Ziegler et al., 2018). These differences in symbiont community are associated with different traits of the diverse Symbiodiniaceae taxa, that might be related to the menthol bleaching tolerance. For instance, taxa with low production of the Photosystem II inihibiting reactive oxygen species (ROS) during menthol exposure, tolerate higher menthol doses than taxa that produce more ROS (Wang et al., 2017). Reactive oxygen species have also been implicated in the breakdown of the symbiosis during heat stress (Helgoe et al., 2024) with large differences in heat stress tolerance between species both within and between Symbiodiniaceae genera (Swain et al., 2017). Identification of symbiont communities in future coral bleaching trials will help elucidate the role of the Symbiodiniaceae community in the menthol bleaching response. In addition, the mechanisms of bleaching and the link between ROS production during menthol exposure and heat stress require further attention.

### Recovery from bleaching

We observed increased minimal fluorescence and tissue color intensity at t2 and t3 compared to t1 after menthol incubation, indicating that chlorophyll and possible symbiont cell density increased in the coral tissue during the recovery phase. This might be related to the retained physiological functionality of the symbionts and suggests a possible symbiont uptake from the environment (Scharfenstein et al., 2022), or proliferation of the remaining symbiont cells in the tissues or of endolithic algae (Galindo-Martínez et al., 2022; Fine et al. 2002). The short time frame for the increase in symbiont density between t1 and t2 (day 2 and day 7 after menthol incubation) suggests that coral fragments may start regaining symbionts within the first week of recovery. This time frame for the onset of recovery is unexpectedly short compared to the observations by Scharfenstein et al. (2022), where symbiont density recovered more slowly. The combined use of menthol with DCMU in that study may explain the difference in recovery times and sets an incentive to systematically explore further bleaching treatments in the future.

Significant decreases in symbiont density with increased menthol exposure observed for *M. digitata*, and *P. cylindrica* could not be fully captured with measurements of minimum chlorophyll fluorescence and tissue color intensity. While these non-invasive proxies enable repeated measures to monitor bleaching progression, they were not sufficiently sensitive to capture the fine-scale differences in symbiont cell density between treatments in our study. Yet overall, the trends across analyses align the successful effect of menthol bleaching.

### Conclusions

Chemical bleaching with menthol effectively produces Symbiodiniaceae-free corals, which can be subsequently used for further investigations, including studies aiming to understand the cellular process involved in the maintenance or breakdown of the coral-algal symbiosis and test the phenotypes of novel associations with exogenous symbionts. This study of menthol bleaching efficiency expanded the toolkit for coral symbiosis research. Based on our data, we recommend adopting bespoke bleaching protocols for different coral species. While these protocols may serve as a starting point for bleaching other coral species, our data showed large species-specific differences, so that the menthol treatment of one species cannot easily be applied to others even closely related species.

Even though the corals were never completely symbiont free, near-complete symbiont depletion of up to 98 % was achieved. Apart from *P. verrucosa*, all coral species survived the menthol treatment without apparent visual impacts, confirming that menthol is a gentle chemical bleaching reagent for stony corals. The underlying reason for the difference in the menthol bleaching response between species remains to be fully elucidated. Future studies of coral bleaching and symbiosis establishment may benefit from this work.

## Acknowledgements

This study was conducted as part of the *Ocean2100* global change simulation project of the Colombian-German Center of Excellence in Marine Sciences (CEMarin) funded by the German Academic Exchange Service. MZ acknowledges funding by the Deutsche Forschungsgemeinschaft (DFG, German Research Foundation) – Project number 469364832 (M.Z.) – SPP 2299/Project number 441832482.

## Data availability

All raw data are provided as supplementary material.

## Author contributions

LB: Investigation, Formal analysis, Writing - Original Draft Preparation. EF: Investigation, Writing – Review & Editing. GP: Investigation, Analysis, Writing – Review & Editing. ALP: Investigation. MR: Investigation. FS: Investigation. MZe: Investigation. MZi: Conceptualization, Resources, Supervision, Writing - Original Draft Preparation, Writing – Review & Editing.

## Supplementary Material

**Fig. S1:**
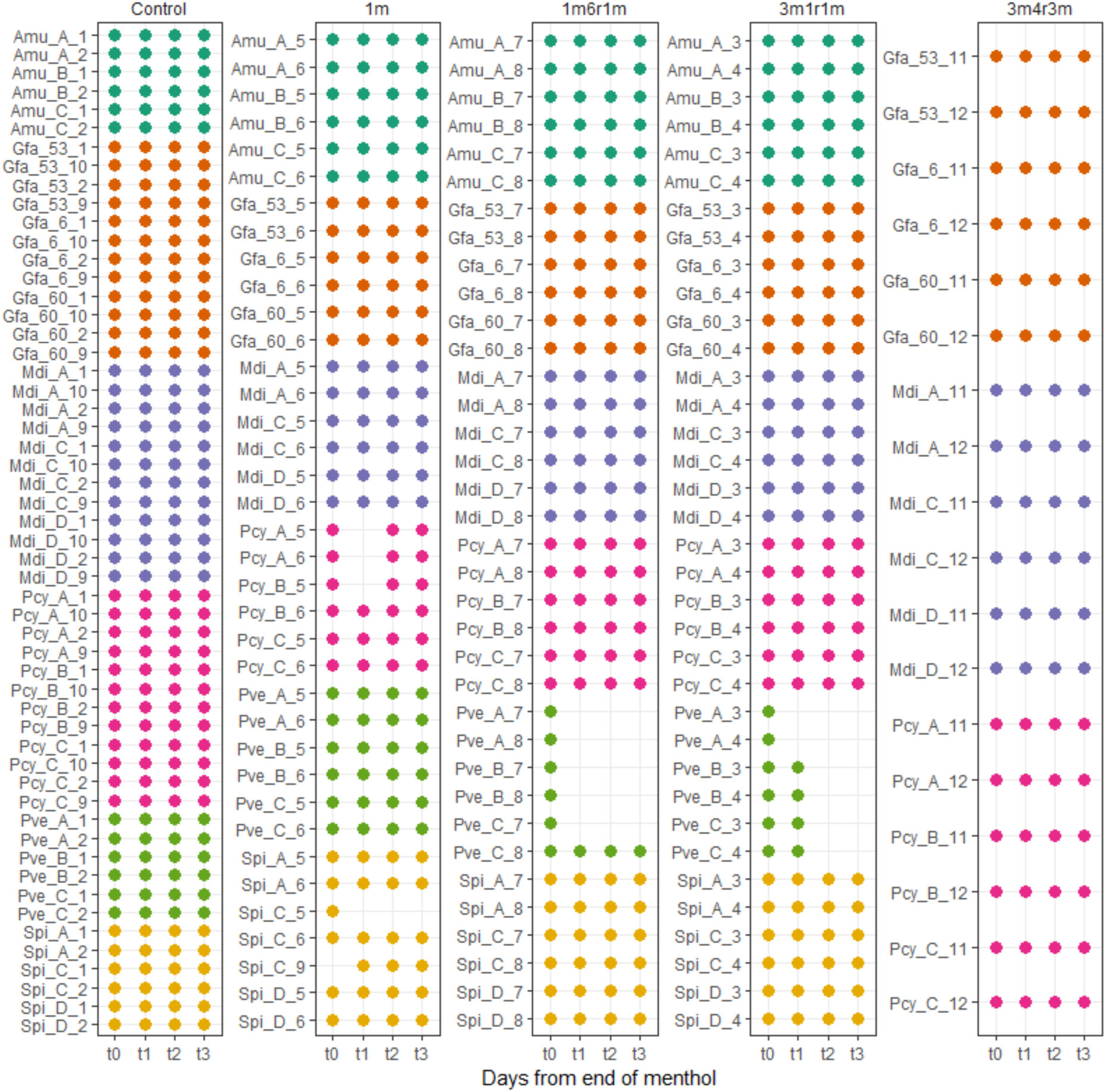
Overview of sampling days: Dots show processed samples by sample-ID (y) at timepoints t0, t1, t2, t3 (x) grouped by menthol treatment (top). Different colors denote different species. The sample-ID is generated following the logic of *A. muricata* = Amu; A, B, C, D = genotype; 1-12 = fragment number (clonal replicates). Missing dots show dead samples or missing measurements. The control consists of more fragments as it was repeated as part of run1 and run2.

**Fig. S2:**
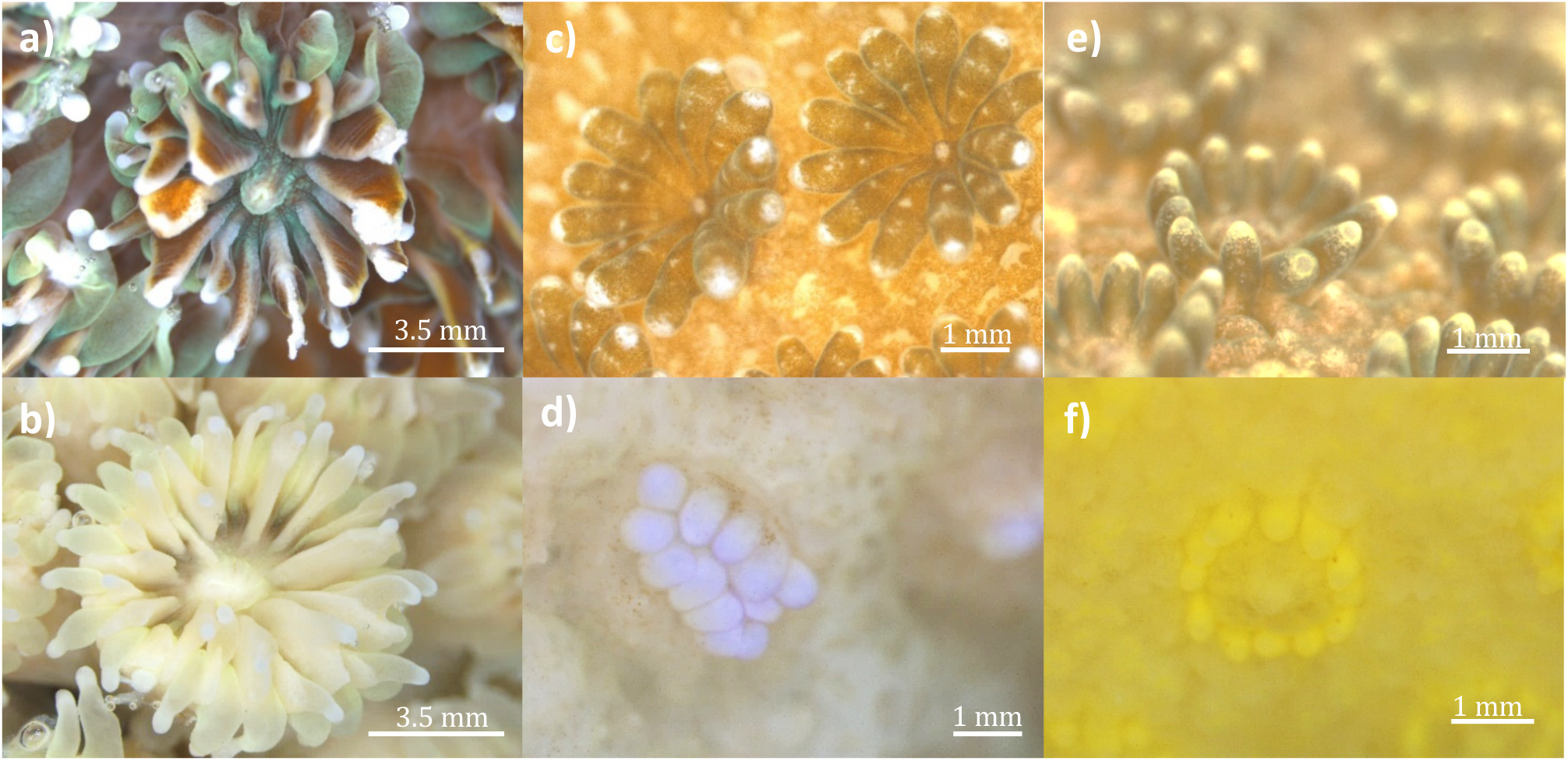
Symbiotic and bleached coral fragments of *G. fascicularis* (a,b), *M. digitata* (c,d), and *P. cylindrica* (e,f) 13 days after menthol treatment. Fragments of the control group (a, c, e) and fragments treated with 0.38 mM menthol in the 3m4r3m protocol (b, d, f) were photographed under a dissecting microscope (Leica MC176 HD).

**Fig. S3:**
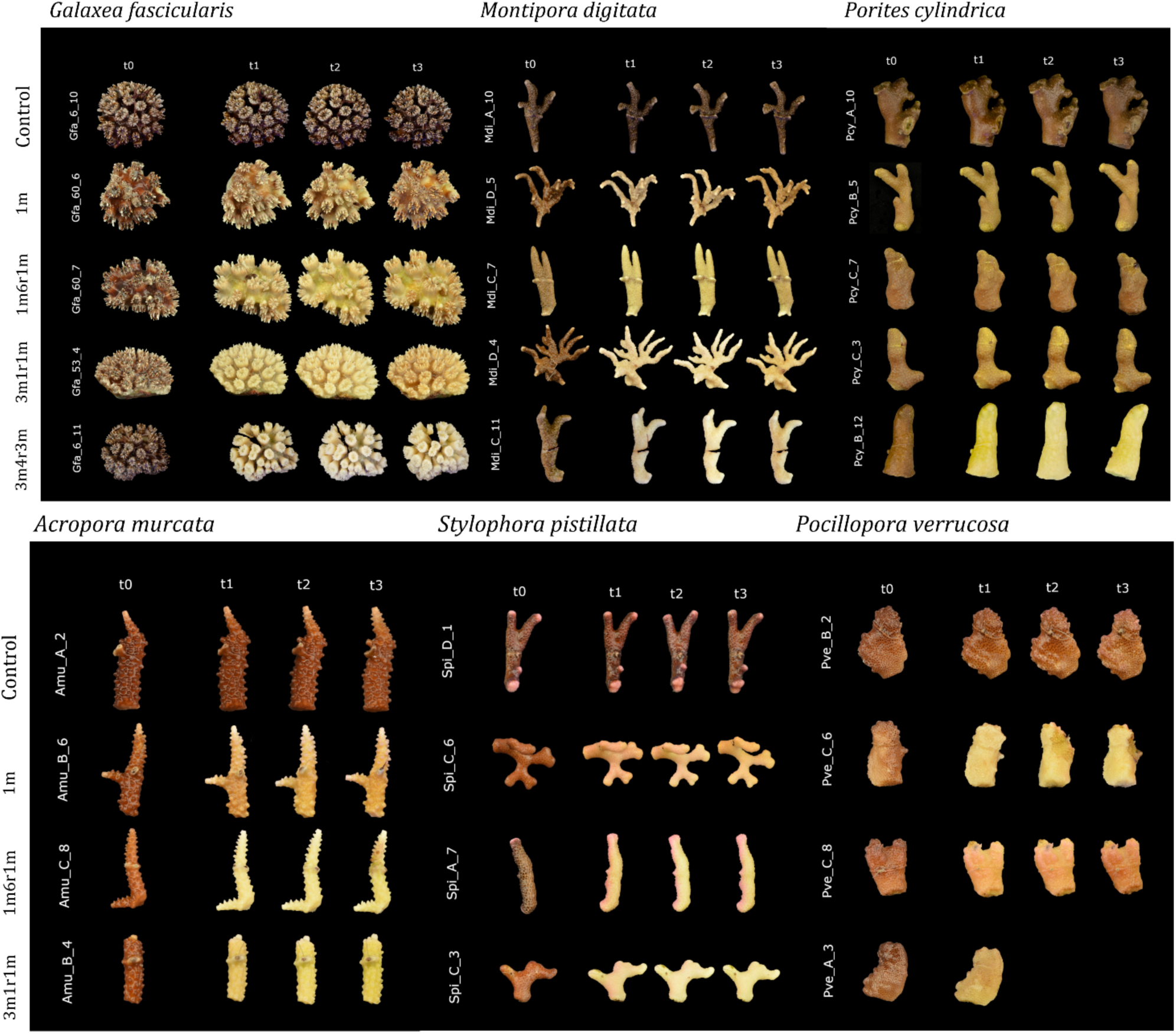
Bleaching series pictures of coral fragments after menthol treatment. Exemplary standardized pictures of *G. fascicularis, S. pistillata, M. digitata* (top) and *A. muricata, S. pistillata,* and *P. verrucosa* (bottom) sorted by timepoint after menthol treatment and menthol treatment. Each row shows the bleaching series in one treatment. The genotype of species varies between menthol treatments (see sample-ID displayed on the left).

**Fig. S4:**
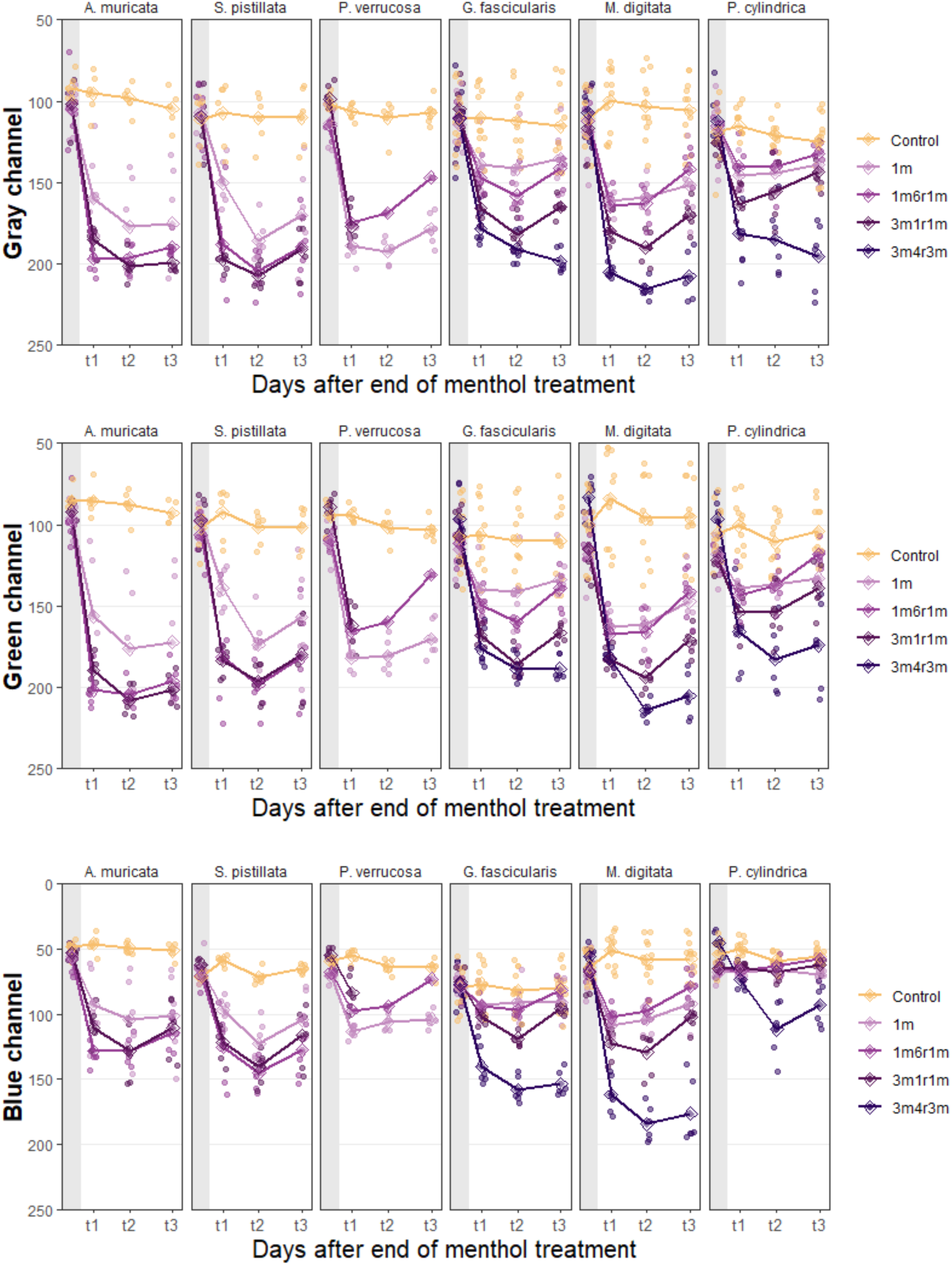
Gray composite, green, and blue channel analysis from standardized pictures (y) per species at the sampling time points t0, t1, t2, t3 of menthol incubation (x). Colored lines show different menthol treatments – darker lines show increased menthol exposure. High values on the y-axis indicate lighter tissue color – low values indicate darker tissue color.

**Tab. S1:**
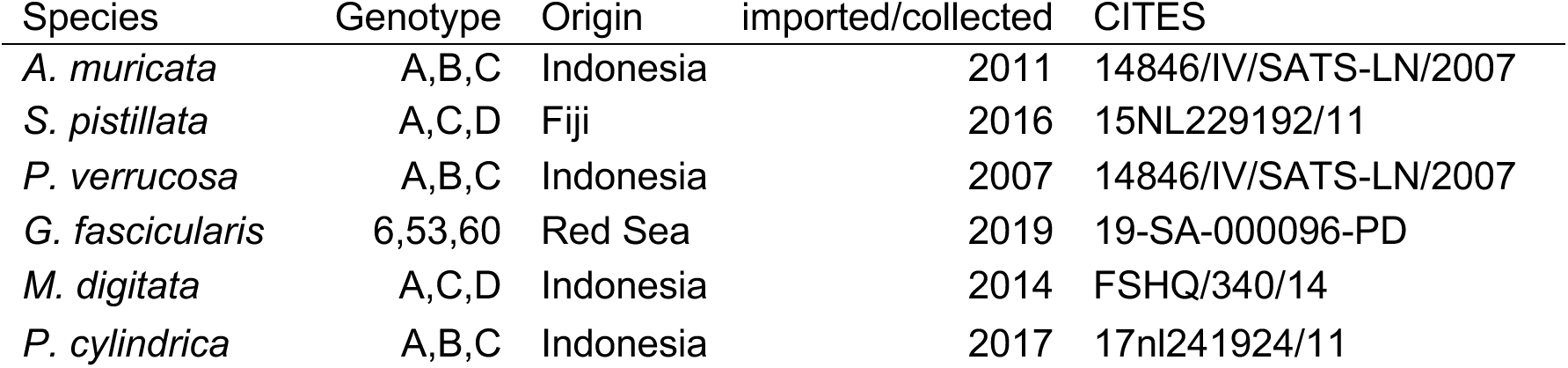
Origin of coral species: Origin, CITES permits and year of acquisition are displayed for the genotypes of six stony coral species.

**Tab. S2:**
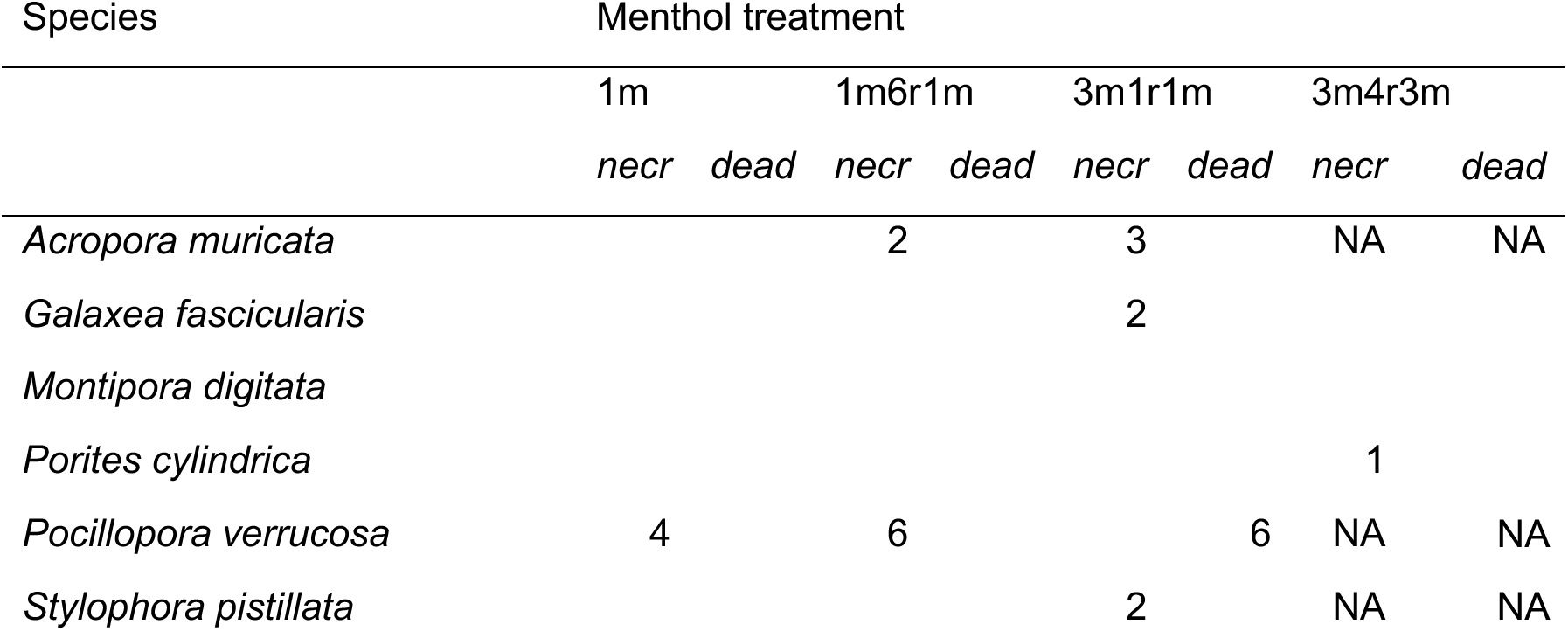
Necrosis (*necr*) and mortality (*dead*) of coral fragments per treatment. Numbers of dead and necrotic samples per species and treatment. Dead fragments also had necrosis. Necrotic fragments were not completely dead. NA = treatment not applied

**Tab. S3:** Repeated measures two-way-ANOVA for minimal fluorescence (F0). The effect of treatment, time (= Day_after_menthol) and their interaction are shown per species. Significant p-values are marked with asterisks (p ≤ 0.05*; p ≤ 0.01**; p ≤ 0.001***)

| Species | Effect | DFn | DFd | F | p |  |
| --- | --- | --- | --- | --- | --- | --- |
| *Acropora muricata* | Treatment | 3 | 15 | 17 | <0.001 | *** |
|  | Day_after_menthol | 3 | 15 | 24 | <0.001 | *** |
|  | Treatment:Day_after_menthol | 9 | 45 | 13 | <0.001 | *** |
| *Galaxea fascicularis* | Treatment | 4 | 20 | 69 | <0.001 | *** |
|  | Day_after_menthol | 1.14 | 5.7 | 124 | <0.001 | *** |
|  | Treatment:Day_after_menthol | 12 | 60 | 32 | <0.001 | *** |
| *Montipora digitata* | Treatment | 4 | 20 | 37 | <0.001 | *** |
|  | Day_after_menthol | 3 | 15 | 69 | <0.001 | *** |
|  | Treatment:Day_after_menthol | 12 | 60 | 22 | <0.001 | *** |
| *Porites cylindrica* | Treatment | 4 | 4 | 3.994 | 0.104 | ns |
|  | Day_after_menthol | 3 | 3 | 15.106 | 0.026 | * |
|  | Treatment:Day_after_menthol | 12 | 12 | 11.24 | <0.001 | *** |
| *Stylophora pistillata* | Treatment | 1.19 | 5.96 | 199 | <0.001 | *** |
|  | Day_after_menthol | 3 | 15 | 70 | <0.001 | *** |
|  | Treatment:Day_after_menthol | 9 | 45 | 30 | <0.001 | *** |

**Tab. S4:**
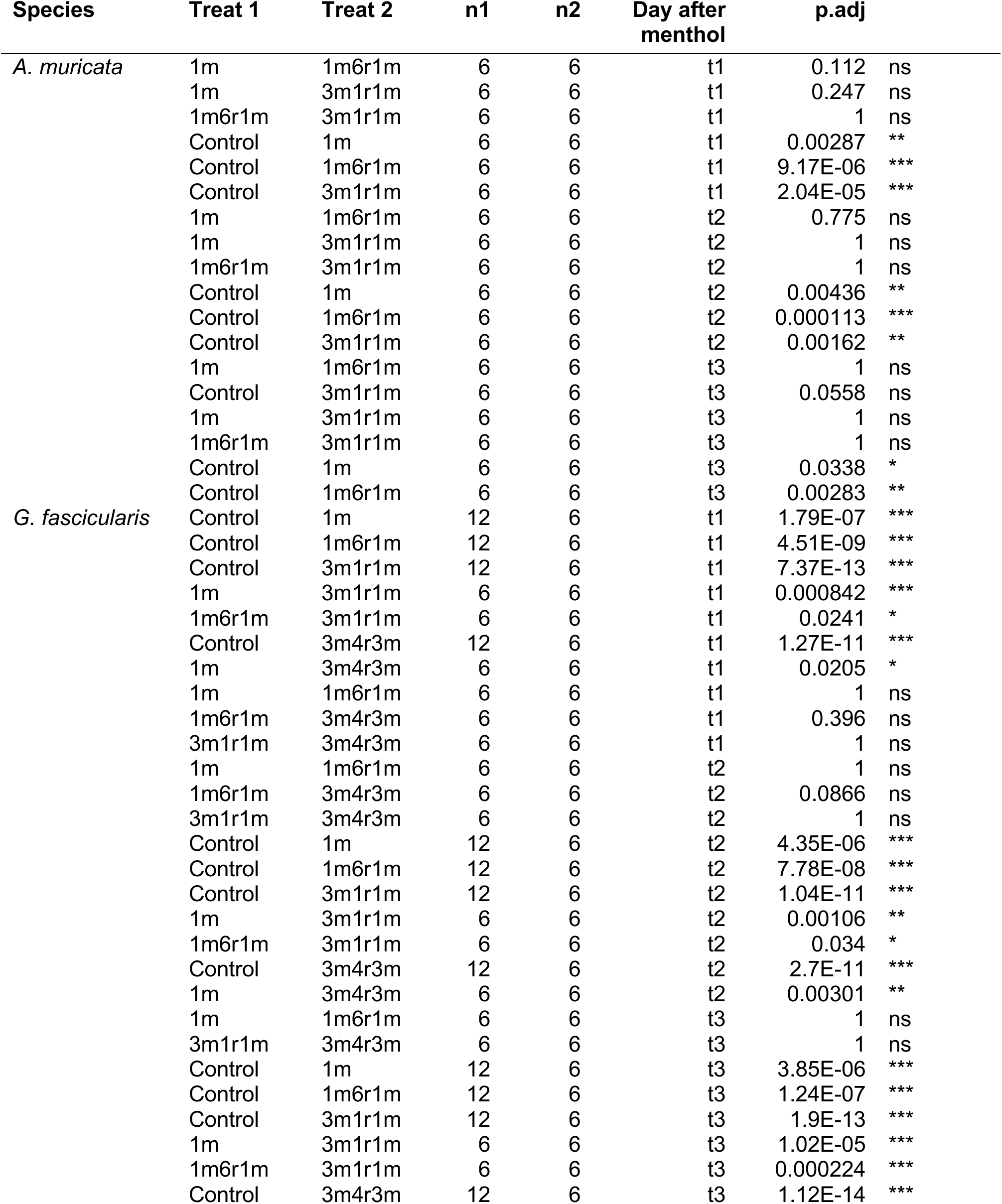
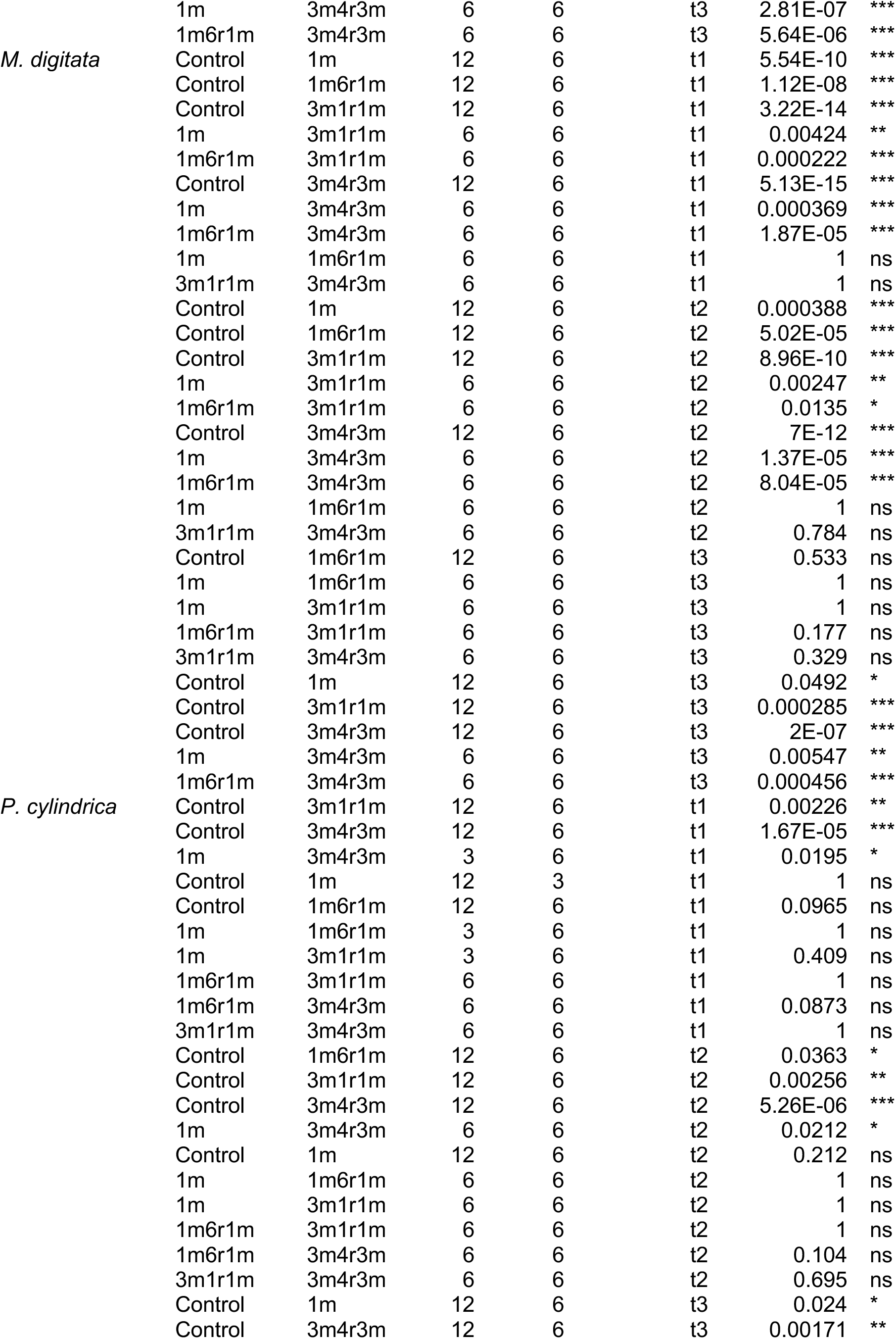
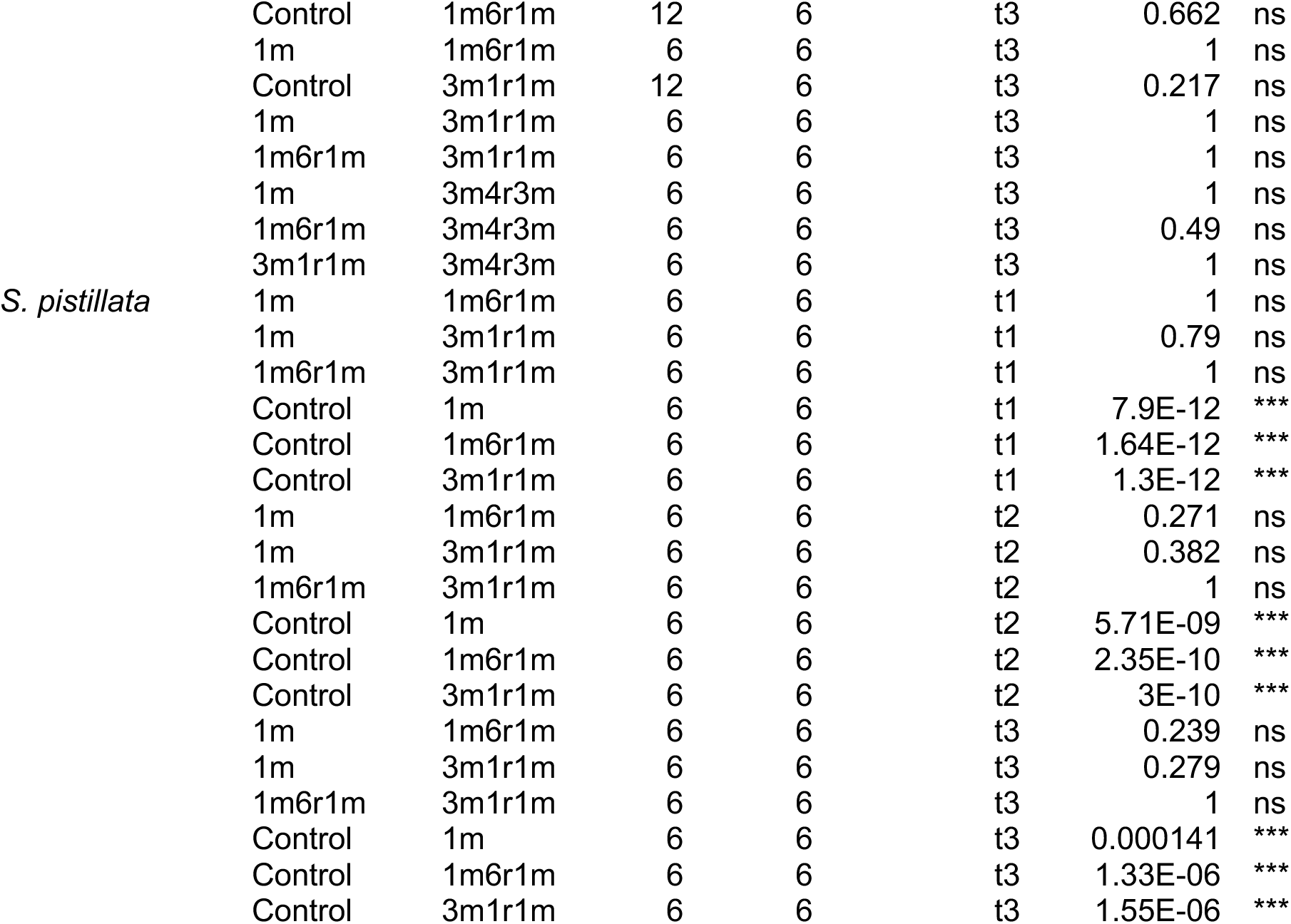
All post-hoc paired t-test values for minimal fluorescence (PAM) data on sampling time points t1, t2, t3 for all menthol treatments and the control. Pairwise comparisons are sorted by species. Asterisks show significant differences, ns = not significant. (p ≤ 0.05*; p ≤ 0.01**; p ≤ 0.001***)

**Tab. S5:**
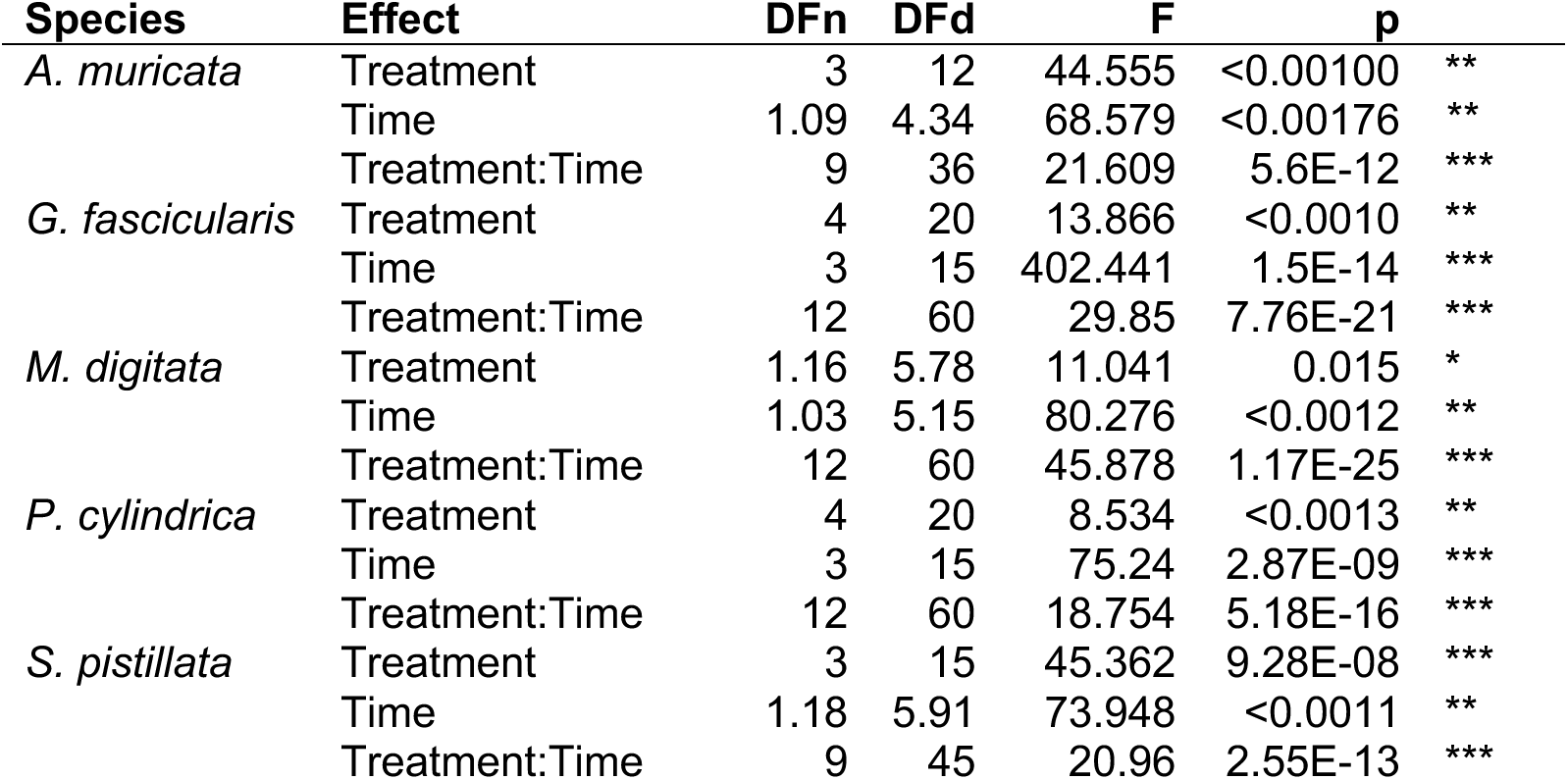
Repeated measures Two-Way-ANOVA table for tissue brightness (red) analysis. The effect of treatment, time (= day after menthol) and their interaction are shown per species. Significant p-values are marked with asterisks (p ≤ 0.05*; p ≤ 0.01**; p ≤ 0.001***)

**Tab. S6:**
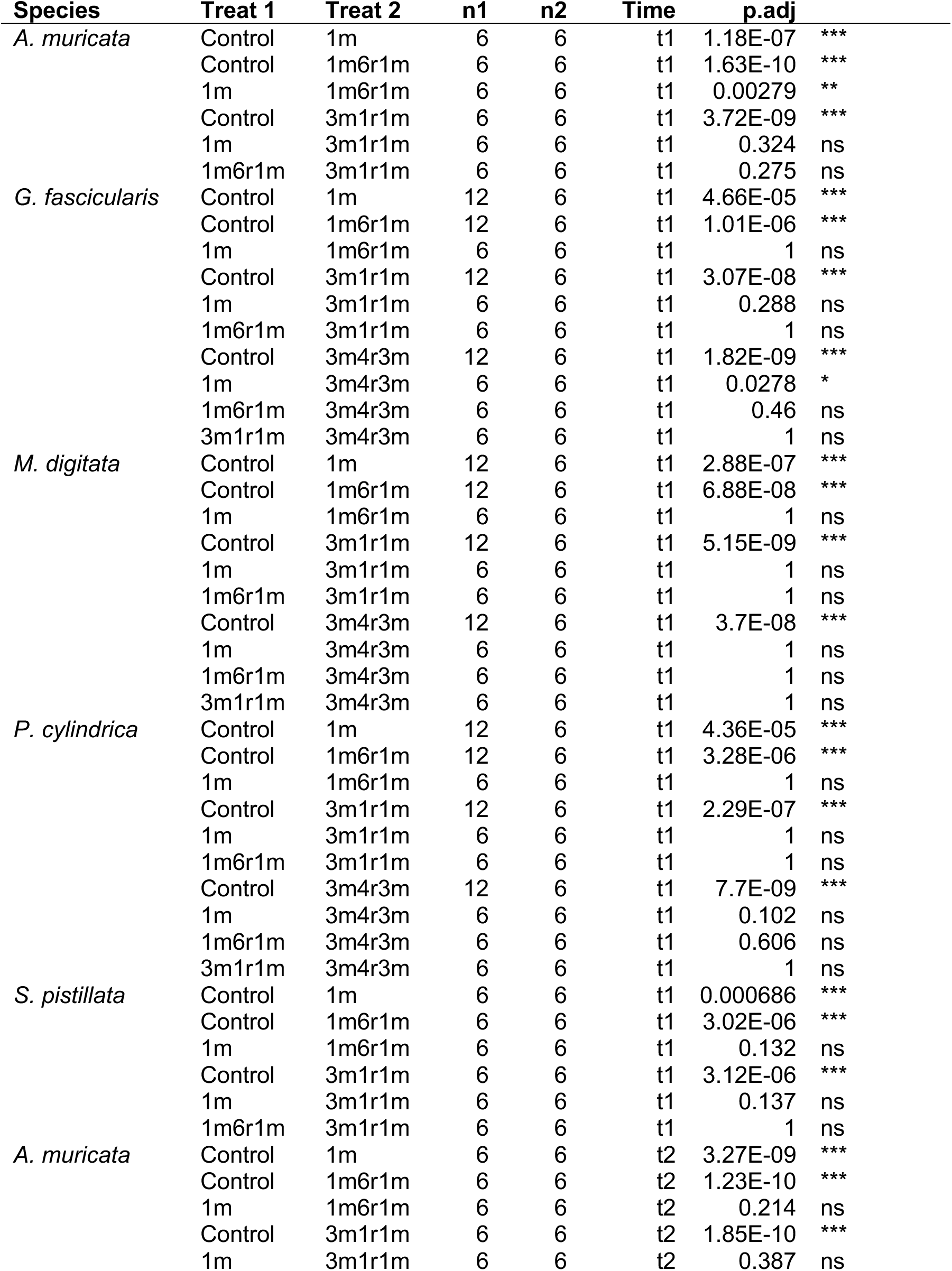
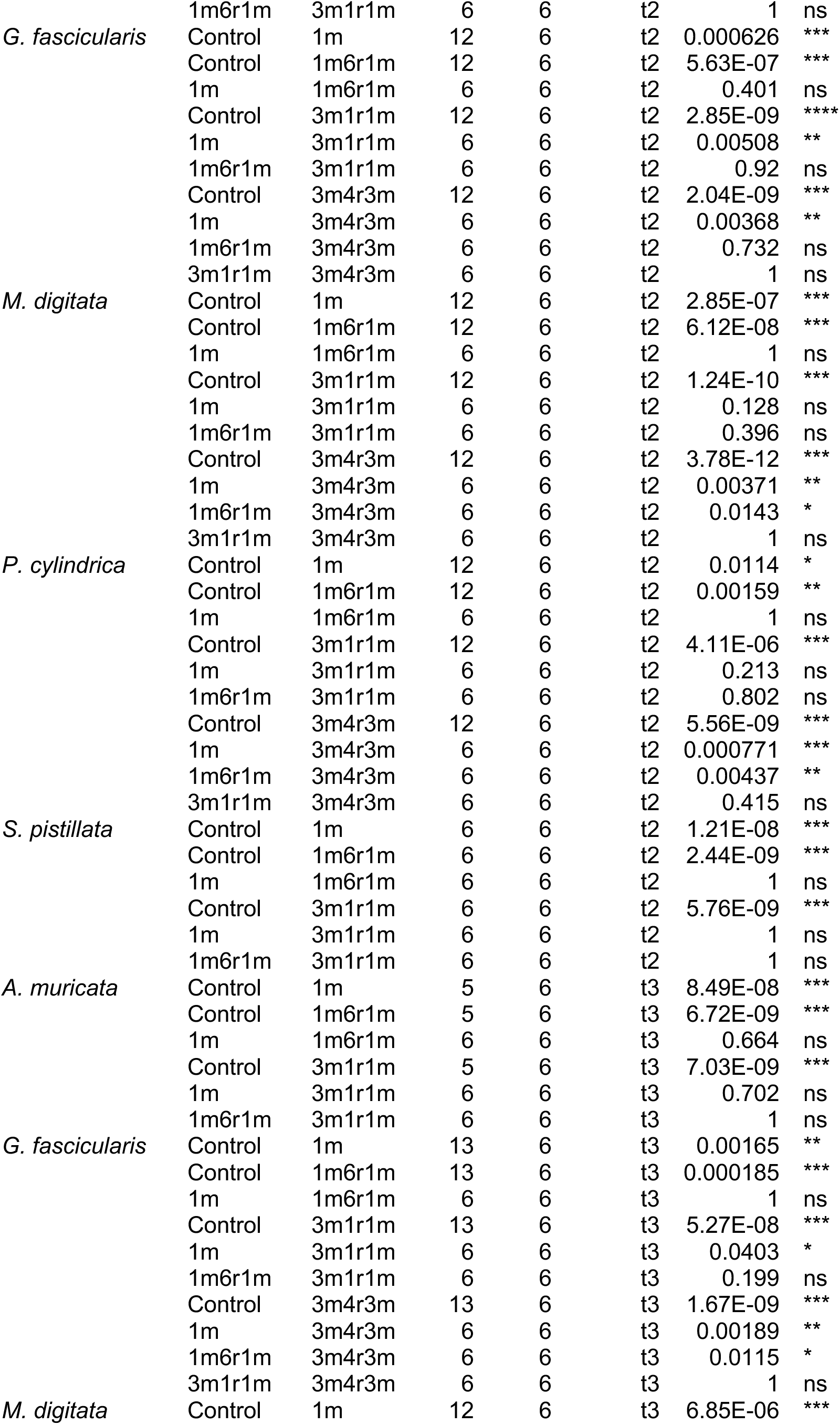
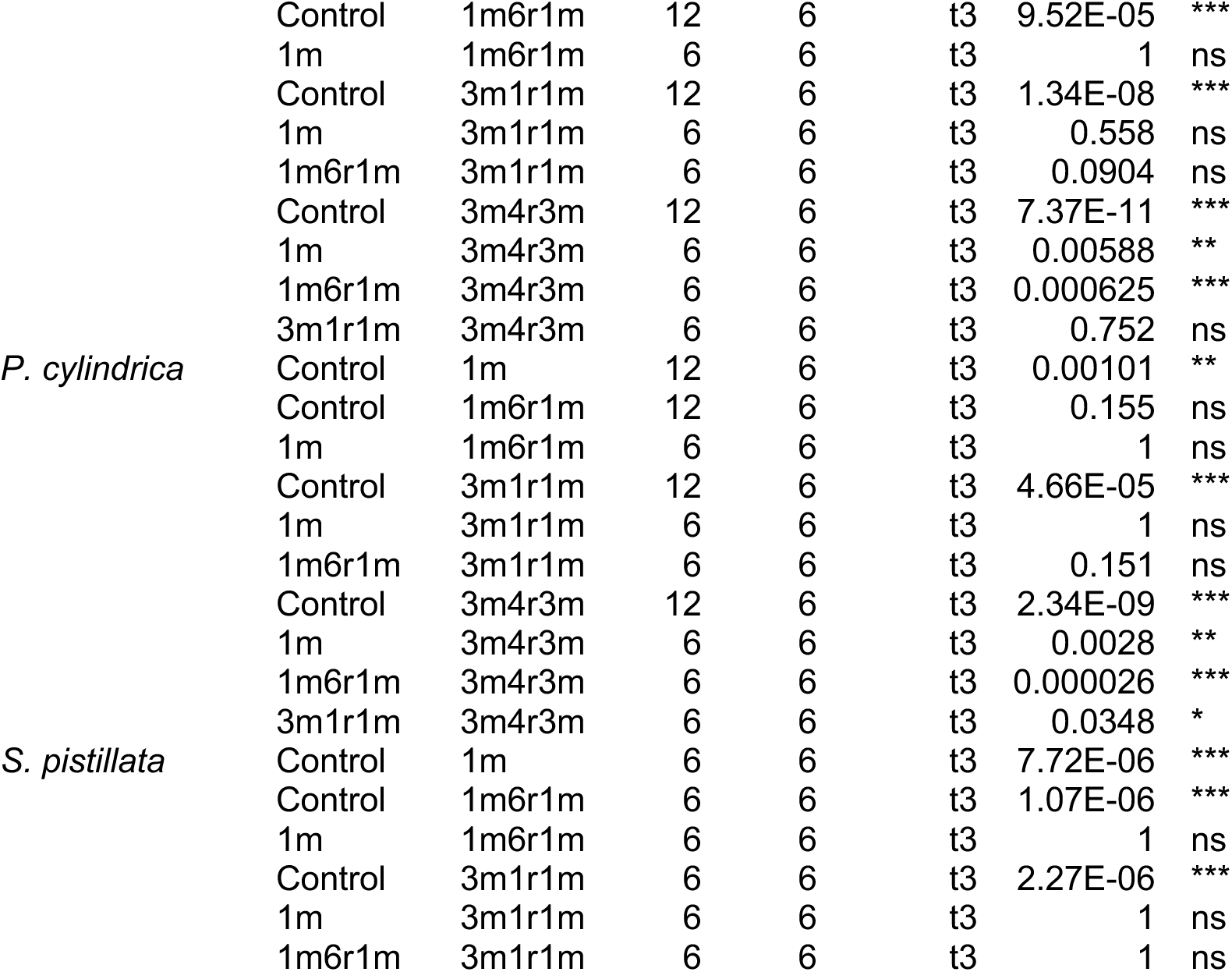
Post-hoc paired t-test values for tissue brightness (red) data on sampling time points t1, t2, t3 for all menthol treatments and the control. Pairwise comparisons are sorted by species. Asterisks show significant differences, ns = not significant. (p ≤ 0.05*; p ≤ 0.01**; p ≤ 0.001***)

**Tab. S7:** Results Turkey-HSD-Test Symbiont density: Pairwise comparison of menthol treatments per species for symbiont density [cells/cm^2^]. Asterisks show significant differences, ns = not significant. (p ≤ 0.05*; p ≤ 0.01**; p ≤ 0.001***)

| Species | Treat 1 | Treat 2 | estimate | conf.low | conf.high | p.adj |  |
| --- | --- | --- | --- | --- | --- | --- | --- |
| *A. muricata* | C | 1m | -1273537 | -1643790 | -903285 | 3.41E-08 | *** |
|  | C | 1m6r1m | -1549343 | -1919596 | -1179090 | 1.2E-09 | *** |
|  | C | 3m1r1m | -1558945 | -1929197 | -1188692 | 1.08E-09 | *** |
|  | 1m | 1m6r1m | -275805 | -646058 | 94447 | 0.192 | ns |
|  | 1m | 3m1r1m | -285407 | -655660 | 84846 | 0.17 | ns |
|  | 1m6r1m | 3m1r1m | -9602 | -379855 | 360651 | 1 | ns |
| *G. fascicularis* | C | 1m | 63307 | -800372 | 926987 | 1 | ns |
|  | C | 1m6r1m | -543905 | -1407584 | 319774 | 0.379 | ns |
|  | C | 3m1r1m | -1131380 | -1995059 | -267701 | 0.00545 | ** |
|  | C | 3m4r3m | -1381432 | -2245112 | -517753 | <0.00156 | *** |
|  | 1m | 1m6r1m | -607213 | -1604503 | 390078 | 0.413 | ns |
|  | 1m | 3m1r1m | -1194687 | -2191978 | -197397 | 0.0126 | * |
|  | 1m | 3m4r3m | -1444740 | -2442031 | -447449 | 0.00186 | ** |
|  | 1m6r1m | 3m1r1m | -587475 | -1584765 | 409816 | 0.446 | ns |
|  | 1m6r1m | 3m4r3m | -837527 | -1834818 | 159764 | 0.134 | ns |
|  | 3m1r1m | 3m4r3m | -250052 | -1247343 | 747238 | 0.949 | ns |
| *M. digitata* | C | 1m | -1030641 | -1842919 | -218363 | 0.00744 | ** |
|  | C | 1m6r1m | -642266 | -1454544 | 170012 | 0.176 | ns |
|  | C | 3m1r1m | -1590339 | -2402617 | -778061 | <0.0010 | *** |
|  | C | 3m4r3m | -2053764 | -2866042 | -1241486 | <0.0010 | *** |
|  | 1m | 1m6r1m | 388375 | -549563 | 1326313 | 0.752 | ns |
|  | 1m | 3m1r1m | -559698 | -1497636 | 378240 | 0.433 | ns |
|  | 1m | 3m4r3m | -1023123 | -1961061 | -85185 | 0.0271 | * |
|  | 1m6r1m | 3m1r1m | -948073 | -1886011 | -10135 | 0.0466 | * |
|  | 1m6r1m | 3m4r3m | -1411498 | -2349436 | -473560 | 0.00119 | ** |
|  | 3m1r1m | 3m4r3m | -463425 | -1401363 | 474513 | 0.613 | ns |
| *P. cylindrica* | C | 1m | -130971 | -842744 | 580803 | 0.983 | ns |
|  | C | 1m6r1m | -318439 | -1030213 | 393334 | 0.696 | ns |
|  | C | 3m1r1m | -639789 | -1351562 | 71984 | 0.0946 | ns |
|  | C | 3m4r3m | -1030837 | -1742610 | -319064 | 0.00186 | ** |
|  | 1m | 1m6r1m | -187469 | -1009354 | 634416 | 0.963 | ns |
|  | 1m | 3m1r1m | -508818 | -1330703 | 313067 | 0.396 | ns |
|  | 1m | 3m4r3m | -899866 | -1721751 | -77981 | 0.0263 | * |
|  | 1m6r1m | 3m1r1m | -321350 | -1143235 | 500535 | 0.789 | ns |
|  | 1m6r1m | 3m4r3m | -712398 | -1534283 | 109487 | 0.115 | ns |
|  | 3m1r1m | 3m4r3m | -391048 | -1212933 | 430837 | 0.646 | ns |
| *P. verrucosa* | C | 1m | -1306655 | -1782134 | -831175 | <0.0010 | *** |
|  | C | 1m6r1m | -1407053 | -1882532 | -931574 | <0.0010 | *** |
|  | C | 3m1r1m | -1374981 | -2264521 | -485440 | 0.00233 | ** |
|  | 1m | 1m6r1m | -100399 | -575878 | 375081 | 0.928 | ns |
|  | 1m | 3m1r1m | -68326 | -957866 | 821214 | 0.996 | ns |
|  | 1m6r1m | 3m1r1m | 32073 | -857468 | 921613 | 1 | ns |
| *S. pistillata* | C | 1m | -1922627 | -2838120 | -1007134 | <0.0011 | *** |
|  | C | 1m6r1m | -2451033 | -3366526 | -1535540 | <0.0010 | *** |
|  | C | 3m1r1m | -2435015 | -3350508 | -1519522 | <0.0010 | *** |
|  | 1m | 1m6r1m | -528406 | -1443899 | 387087 | 0.393 | ns |
|  | 1m | 3m1r1m | -512388 | -1427881 | 403105 | 0.419 | ns |
|  | 1m6r1m | 3m1r1m | 16018 | -899475 | 931511 | 1 | ns |

## References

Berkelmans, R., & Van Oppen, M. J. H. (2006). The role of zooxanthellae in the thermal tolerance of corals: A “nugget of hope” for coral reefs in an era of climate change. Proceedings of the Royal Society B: Biological Sciences, 273(1599), 2305–2312. 10.1098/rspb.2006.3567

Buerger, P., Alvarez-Roa, C., Coppin, C. W., Pearce, S. L., Chakravarti, L. J., Oakeshott, J. G., Edwards, O. R., & Van Oppen, M. J. H. (2020). M A R I N E B I O L O G Y Heat-evolved microalgal symbionts increase coral bleaching tolerance. https://www.science.org

Chan, W.Y., Meyers, L., Rudd, D., Topa, S.H., van Oppen, M.J.H. (2023). Heat-evolved algal symbionts enhance bleaching tolerance of adult corals without trade-off against growth. Global Change Biology, 29, 6945–6968. 10.1111/gcb.16987

Cignoni, P., Callieri, M., Corsini, M., Dellepiane, M., Ganovelli, F., & Ranzuglia, G. (2008). MeshLab: an Open-Source Mesh Processing Tool.

Coffroth, M. A., Poland, D. M., Petrou, E. L., Brazeau, D. A., & Holmberg, J. C. (2010). Environmental symbiont acquisition may not be the solution to warming seas for reef-building corals. PLoS ONE, 5(10). 10.1371/journal.pone.0013258

Darling, E. S., Alvarez-Filip, L., Oliver, T. A., McClanahan, T. R., & Côté, I. M. (2012). Evaluating life-history strategies of reef corals from species traits. Ecology Letters, 15(12), 1378–1386. 10.1111/j.1461-0248.2012.01861.x

Eddy TD, Lam VWY, Reygondeau G, Cisneros-Montemayor AM, Greer K, Palomares MLD, Bruno JF, Ota Y, Cheung WWL (2021) Global decline in capacity of coral reefs to provide ecosystem services. One Earth 4:1278–1285. 10.1016/j.oneear.2021.08.016

Ferrara, E.F., Bauer, L., Puntin, G., Bautz, F., Celayir, S., Do, M.-S., Eck, F., Heider, M., Wissel, P., Arnold, A., Wilke, T., Reichert, J., Ziegler, M. (2024). RGB color indices as proxy for symbiont cell density and chlorophyll content during coral bleaching. bioRxiv 2024.12.20.629333. 10.1101/2024.12.20.629333

Fine, M., & Loya, Y. (2002). Endolithic algae: an alternative source of photoassimilates during coral bleaching. Proceedings of the Royal Society of London. Series B: Biological Sciences, 269(1497), 1205–1210. 10.1098/rspb.2002.1983

Frankowiak, K., Wang, X. T., Sigman, D. M., Gothmann, A. M., Kitahara, M. V, Mazur, M., Meibom, A., & Stolarski, J. (2016). P A L E O N T O L O G Y Photosymbiosis and the expansion of shallow-water corals.

Frölicher, T. L., Fischer, E. M., & Gruber, N. (2018). Marine heatwaves under global warming. Nature, 560(7718), 360–364. 10.1038/s41586-018-0383-9

Galindo-Martínez, C. T., Weber, M., Avila-Magaña, V., Enríquez, S., Kitano, H., Medina, M., & Iglesias-Prieto, R. (2022). The role of the endolithic alga Ostreobium spp. during coral bleaching recovery. Scientific reports, 12(1), 2977. 10.1038/s41598-022-07017-6

H. Wickham. ggplot2: Elegant Graphics for Data Analysis. Springer-Verlag New York, 2016. https://ggplot2.tidyverse.org v. 3.4.4

Helgoe, J., Davy, S. K., Weis, V. M., & Rodriguez-Lanetty, M. (2024). Triggers, cascades, and endpoints: Connecting the dots of coral bleaching mechanisms. Biological Reviews, 99(3), 715–752. 10.1111/brv.13042

Hoegh-Guldberg, O., Mumby, P. J., Hooten, A. J., Steneck, R. S., Greenfield, P., Gomez, E., Harvell, C. D., Sale, P. F., Edwards, A. J., Caldeira, K., Knowlton, N., Eakin, C. M., Iglesias-Prieto, R., Muthiga, N., Bradbury, R. H., Dubi, A., & Hatziolos, M. E. (2007). Coral Reefs Under Rapid Climate Change and Ocean Acidification. https://www.science.org

Hughes, T. P., Kerry, J. T., Baird, A. H., Connolly, S. R., Dietzel, A., Eakin, C. M., Heron, S. F., Hoey, A. S., Hoogenboom, M. O., Liu, G., McWilliam, M. J., Pears, R. J., Pratchett, M. S., Skirving, W. J., Stella, J. S., & Torda, G. (2018). Global warming transforms coral reef assemblages. Nature, 556(7702), 492–496. 10.1038/s41586-018-0041-2

IPCC, 2023: Climate Change 2023: Synthesis Report. Contribution of Working Groups I, II and III to the Sixth Assessment Report of the Intergovernmental Panel on Climate Change [Core Writing Team, H. Lee and J. Romero (eds.)]. IPCC, Geneva, Switzerland, pp. 35–115, doi: 10.59327/IPCC/AR6-9789291691647

Kassambara A (2023). rstatix: Pipe-Friendly Framework for Basic Statistical Tests. R package version 0.7.2, https://rpkgs.datanovia.com/rstatix/.

Kirillov, A., Mintun, E., Ravi, N., Mao, H., Rolland, C., Gustafson, L., Xiao, T., Whitehead, S., Berg, A. C., Lo, W.-Y., Dollár, P., & Girshick, R. (2023). Segment Anything. ArXiv*:2304.02643*. http://arxiv.org/abs/2304.02643

Madin, J. S., Anderson, K. D., Andreasen, M. H., Bridge, T. C. L., Cairns, S. D., Connolly, S. R., Darling, E. S., Diaz, M., Falster, D. S., Franklin, E. C., Gates, R. D., Hoogenboom, M. O., Huang, D., Keith, S. A., Kosnik, M. A., Kuo, C. Y., Lough, J. M., Lovelock, C. E., Luiz, O., … Baird, A. H. (2016). The Coral Trait Database, a curated database of trait information for coral species from the global oceans. Scientific Data, 3. 10.1038/sdata.2016.17

Matthews, J. L., S. A. E., O. C. A., G. A. R., W. V. M., D. S. K. (2016). Menthol-induced bleaching rapidly and effectively provides experimental aposymbiotic sea anemones (Aiptasia sp.) for symbiosis investigations. Journal of Experimental Biology, 219, 306–310. 10.1242/jeb.128934

McLachlan, R. H., Price, J. T., Solomon, S. L., & Grottoli, A. G. (2020). Thirty years of coral heat-stress experiments: a review of methods. In Coral Reefs (Vol. 39, Issue 4, pp. 885–902). Springer Science and Business Media Deutschland GmbH. 10.1007/s00338-020-01931-9

Meron, D., Maor-landaw, K., Eyal, G., Elifantz, H., Banin, E., Loya, Y., & Levy, O. (2020). The complexity of the holobiont in the red sea coral Euphyllia paradivisa under heat stress. Microorganisms, 8(3). 10.3390/microorganisms8030372

Muscatine, L., Falkowski, P. G., Porter, J. W., & Dubinsky, Z. (1984). Fate of Photosynthetic Fixed Carbon in Light- and Shade-Adapted Colonies of the Symbiotic Coral Stylophora pistillata (Vol. 222, Issue 1227). https://about.jstor.org/terms

Puntin, G., Craggs, J., Hayden, R., Engelhardt, K. E., McIlroy, S., Sweet, M., Baker, D. M., & Ziegler, M. (2023). The reef-building coral Galaxea fascicularis: a new model system for coral symbiosis research. Coral Reefs, 42(1), 239–252. 10.1007/s00338-022-02334-8

R Core Team (2021). R: A language and environment for statistical computing. R Foundation for Statistical Computing, Vienna, Austria.URL https://www.R-project.org/.

Rades, M., Schubert, P., Ziegler, M., Kröckel, M., & Reichert, J. (2022). Building plan for a temperature-controlled multi-point stirring incubator. protocols.io. 10.17504/protocols.io.dm6gpb34dlzp/v1

Reichert, J., Tirpitz, V., Anand, R., Bach, K., Knopp, J., Schubert, P., Wilke, T., & Ziegler, M. (2021). Interactive effects of microplastic pollution and heat stress on reef-building corals. Environmental Pollution, 290(July), 118010. 10.1016/j.envpol.2021.118010

Scharfenstein, H. J., Chan, W. Y., Buerger, P., Humphrey, C., & van Oppen, M. J. H. (2022). Evidence for de novo acquisition of microalgal symbionts by bleached adult corals. ISME Journal, 16(6), 1676–1679. 10.1038/s41396-022-01203-0

Schubert, P., & Wilke, T. (2018). Coral Microcosms: Challenges and Opportunities for Global Change Biology. In Corals in a Changing World (pp. 143–175). InTech. 10.5772/intechopen.68770

Silverstein, R. N., Cunning, R., & Baker, A. C. (2015). Change in algal symbiont communities after bleaching, not prior heat exposure, increases heat tolerance of reef corals. Global Change Biology, 21(1), 236–249. 10.1111/gcb.12706

Souter, D., Planes, S., Wicquart, J., Logan, M., Obura, D., & Staub, F. (2020). Status of Coral Reefs of the World: 2020 Summary for Policymakers Summary for Policymakers-Status of Coral Reefs of the World: 2020 Value of coral reefs.

Steen, R. G., & Muscatine, L. (1987). Low Temperature Evokes Rapid Exocytosis of Symbiotic Algae by a Sea Anemone. In Bulletin (Vol. 172, Issue 2). https://about.jstor.org/terms

Steneck, R. S., & Torres, R. (2023). Trends in Dominican Republic Coral Reef Biodiversity 2015–2022. Diversity, 15(3), 389. 10.3390/d15030389

Strader, M. E., & Quigley, K. M. (2022). The role of gene expression and symbiosis in reef-building coral acquired heat tolerance. Nature Communications, 13(1). 10.1038/s41467-022-32217-z

Swain, T. D., Chandler, J., Backman, V., & Marcelino, L. (2017). Consensus thermotolerance ranking for 110 Symbiodinium phylotypes: An exemplar utilization of a novel iterative partial-rank aggregation tool with broad application potential. Functional Ecology, 31(1), 172–183. 10.1111/1365-2435.12694

Van Hooidonk, R., Maynard, J., Tamelander, J., Gove, J., Ahmadia, G., Raymundo, L., Williams, G., Heron, S. F., & Planes, S. (2016). Local-scale projections of coral reef futures and implications of the Paris Agreement. Scientific Reports, 6. 10.1038/srep39666

Voolstra, C. R., Buitrago-López, C., Perna, G., Cárdenas, A., Hume, B. C. C., Rädecker, N., & Barshis, D. J. (2020). Standardized short-term acute heat stress assays resolve historical differences in coral thermotolerance across microhabitat reef sites. Global Change Biology, 26(8), 4328–4343. 10.1111/gcb.15148

Wang, J. T., Keshavmurthy, S., Chu, T. Y., & Chen, C. A. (2017). Diverse responses of Symbiodinium types to menthol and DCMU treatment. PeerJ, 2017(10). 10.7717/peerj.3843

Wang, J.-T., Chen, Y.-Y., Tew, K. S., Meng, P.-J., & Chen, C. A. (2012a). Physiological and biochemical performances of menthol-induced aposymbiotic corals. PLoS ONE, 7(9), e46406. 10.1371/journal.pone.0046406

Zamoum, T., & Furla, P. (2012). Symbiodinium isolation by NaOH treatment. Journal of Experimental Biology, 215(22), 3875–3880. 10.1242/jeb.074955

Ziegler, M., Eguíluz, V. M., Duarte, C. M., & Voolstra, C. R. (2018). Rare symbionts may contribute to the resilience of coral-algal assemblages. ISME Journal, 12(1), 161–172. 10.1038/ismej.2017.151

